# Joint changes in RNA, RNA polymerase II, and promoter activity through the cell cycle identify non-coding RNAs involved in proliferation

**DOI:** 10.1101/2021.02.12.430890

**Authors:** Siv Anita Hegre, Helle Samdal, Antonin Klima, Endre B. Stovner, Kristin G. Nørsett, Nina Beate Liabakk, Lene Christin Olsen, Konika Chawla, Per Arne Aas, Pål Sætrom

## Abstract

Proper regulation of the cell cycle is necessary for normal growth and development of all organisms. Conversely, altered cell cycle regulation often underlies proliferative diseases such as cancer. Long non-coding RNAs (lncRNAs) are recognized as important regulators of gene expression and are often found dysregulated in diseases, including cancers. However, identifying lncRNAs with cell cycle functions is challenging due to their often low and cell-type specific expression. We present a highly effective method that analyses changes in promoter activity, transcription, and RNA levels for identifying genes enriched for cell cycle functions. Specifically, by combining RNA sequencing with ChIP sequencing through the cell cycle of synchronized human keratinocytes, we identified 1009 genes with cell cycle-dependent expression and correlated changes in RNA polymerase II occupancy or promoter activity as measured by histone 3 lysine 4 trimethylation (H3K4me3). These genes were highly enriched for genes with known cell cycle functions and included 59 lncRNAs. We selected four of these lncRNAs – *AC005682.5, RP11-132A1.4, ZFAS1*, and *EPB41L4A-AS1* – for further experimental validation and found that knockdown of each of the four lncRNAs affected cell cycle phase distributions and reduced proliferation in multiple cell lines. These results show that many genes with cell cycle functions have concomitant cell-cycle dependent changes in promoter activity, transcription, and RNA levels and support that our multi-omics method is well suited for identifying lncRNAs involved in the cell cycle.

## Introduction

Genome-wide gene expression studies have revealed that several genes are regulated in a cell cycle-specific manner [1–5]. Many of these genes are involved in basic cellular processes, such as cell cycle control, DNA repair, DNA replication, and chromosome segregation [2, 6]. One example is the cyclins, which in complex with cyclin-dependent kinases (CDKs), control cell cycle progression. The cyclins are periodically expressed throughout the cell cycle; the E-type cyclins *CCNE1* and *CCNE2* are G1/S-specific [2, 7, 8], whereas the B-type cyclins *CCNB1* and *CCNB2* are G2/M-specific [2, 9, 10].

Long non-coding RNAs (lncRNAs) have emerged as important regulators of gene expression at the epigenetic, transcriptional, and translational level, and are recognized as key modulators in several cancers as well as neurological, autoimmune, and cardiovascular diseases. LncRNAs are more than 200 nucleotides long with little or no protein coding potential, and they generally have a more cell type-specific expression pattern compared to mRNAs [11]. LncRNAs are classified into five main categories according to where they are encoded in the genome in relation to mRNAs: sense, antisense, bi-directional, intergenic, and intronic. They are able to regulate the gene expression at the transcriptional level by acting as signals, guides, scaffolds, or decoys [12]. Most lncRNAs are transcribed by RNA Polymerase II (Pol II) and are poly-adenylated and 5’-capped like mRNAs [13]. A curated knowledgebase of lncRNAs from existing databases and published literature indicates that there are more than 268 000 human lncRNA transcripts, and only a few of them have known functional roles [14].

Several lncRNAs are involved in the cell cycle, possibly through the regulation of other well-known cell cycle regulators like the cyclins, p53, retinoblastoma protein (RB), CDKs, and the CDK inhibitors [15]. A known cell cycle-associated lncRNA is the growth arrest-specific transcript 5 (*GAS5*), which is found downregulated in several cancers where its overexpression results in cell cycle arrest or apoptosis. In prostate cancer *GAS5* functions as a tumor suppressor that inhibits proliferation by targeting the CDK inhibitor p27 [16]. Another known cell cycle-associated lncRNA is the zinc finger antisense 1 (*ZFAS1*). *ZFAS1* can act as an oncogene in some cancer types [17, 18], and as a tumor suppressor in others [19], possibly depending on both type of tissue and state of progression.

The function of the majority of lncRNAs is still unknown, as only about 500-1500 have been functionally characterized. As RNA molecules, lncRNAs need a physical proximity to exert their function. Thus, the subcellular localization of lncRNAs provides important information regarding their potential function [20]. For example, nuclear enriched lncRNAs can act as transcriptional and epigenetic regulators, but they are unlikely to have any coding potential since translation occurs in the cytoplasm. In general, most lncRNAs demonstrate a stronger nuclear localization than mRNAs [21]. Moreover, a higher cell type specificity means that targeting lncRNAs supposedly have less side-effects than targeting protein coding genes [22].

Some lncRNAs can regulate gene transcription by modulating histone modifications [23], although little is known about how lncRNAs are transcriptionally regulated [24]. Histone modifications such as H3K4me3 and H3K27me3 are considered key epigenetic regulators of transcription. H3K4me3 is a mark of actively transcribed genes and H3K27me3 is associated with silenced genes [25]. Previous studies have demonstrated that RNA sequencing (RNA-seq) combined with chromatin immunoprecipitation sequencing (ChIP-seq) are useful for detecting transcriptional fluctuations by correlating gene expression with changes in histone modifications [26, 27]. A study from Wan et.al combined RNA-seq with ChIP-seq and identified differential peaks for H3K4me3 and H3K27me3 around the promoter area and at enhancer regions of differentially expressed lncRNAs in an Alzheimer’s disease mouse model compared to control, suggesting that most of these lncRNA genes were transcriptionally regulated by histone modifications [28]. Since the majority of lncRNAs are spatially and temporally regulated and expressed, ChIP-seq is a sensitive method for capturing these changes by identifying enriched peak regions of histone modifications and other transcriptional regulators [24].

Building on our previous work, which identified protein coding genes with tissue-specific cell cycle-dependent expression [1], we set out to identify lncRNAs with cell cycle-dependent expression and potential cell cycle functions. By combining RNA-seq and Pol II, H3K4me3, and H3K27me3 ChIP-seq data from synchronized HaCaT cells, we identified genes where expression and ChIP-seq signal were correlated and varied dependent on the cell cycle. Genes with high correlation to Pol II and H3K4me3 were strongly enriched for cell cycle functions. From the RNA-seq data we identified 99 lncRNAs with cell cycle-dependent expression profiles; 48, 31, and 15 of these had highly correlated Pol II, H3K4me3, or H3K27me3 signals, respectively. We selected four lncRNAs for further functional characterization and showed that knockdown of these lncRNAs affected cell cycle phase distributions and reduced cell proliferation in multiple cell lines.

## Results

### Total RNA sequencing of HaCaT cells identifies cell cycle genes

Our group previously published a microarray-based study of cell cycle synchronized HaCaT cells identifying a set of genes with cell cycle-dependent expression and strong enrichment for known cell cycle functions [1]. Although our microarray-based study identified several genes with significant periodic expression patterns during the cell cycle, the microarrays precluded the detection of non-coding RNA (ncRNA) transcripts. To identify ncRNAs that are differentially expressed during cell cycle, we therefore set out to do total RNA-seq on synchronized HaCaT cells.

To study cell cycle regulated genes, it is essential to obtain a proper cell synchronization. In two independent experiments (Epi1, Epi2), we used a double thymidine block to arrest and subsequently release HaCaT cells at the G1/S transition, and collected cells every third hour for 24 hours, covering approximately two cycles of DNA replication (Figure 1a-b). Flow cytometry analyses of the cells’ DNA content showed that the distributions of cells through the cell cycle were reproducible between the two independent experiments (Figure 1a). Upon release from thymidine block, approximately 90% of the cells progressed into S phase and continued through the cell cycle (Figure 1b). Cells gradually lost synchrony, such that 70% instead of 90% of the cells synchronously re-entered the second S phase. Based on these results, we considered the HaCaT cells to be effectively synchronized and proceeded with gene expression profiling of the two experiments by using total RNA-seq.

**Fig. 1.**
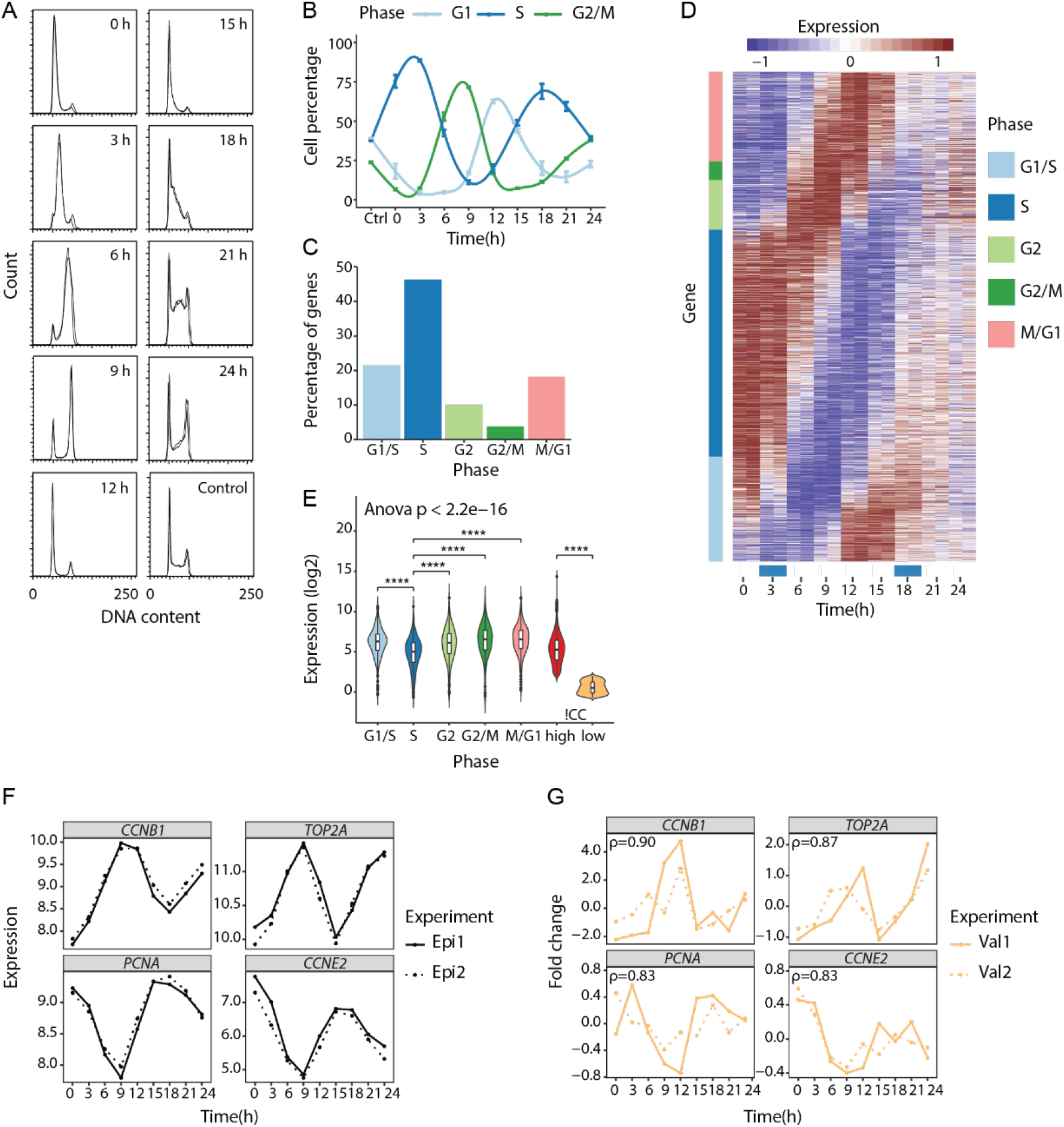
Total RNA-seq of synchronized HaCaT cells identifies cyclic gene expression patterns. **a** HaCaT cells were synchronized by double thymidine block in two independent experiments (Epi1, Epi2) and cell synchrony was monitored by flow cytometry of propidium iodide-stained cells. The figure shows superimposed DNA content profiles of the two replicate experiments for each time point. Horizontal axes show DNA content (arbitrary units) and vertical axes show the number of cells with the corresponding DNA content. Control is unsynchronized cells. **b** Percentage of cells assigned to G1, S, and G2/M phases for each of the time points analyzed. Values and error bars are averages and standard deviations (n = 2). **c** Percentage of cell cycle genes assigned to G1/S (21.5%), S (46.3%), G2 (10.1%), G2/M (3.8%), and M/G1 (18.2%) phases. **d** Heatmap showing the expression changes of the cell cycle genes relative to their median expression. Colour bars in the gene margin (y axis) show the genes’ assigned cell cycle phase; blue bars above the time points (x axis) show the time points having the highest percentage of S phase cells. **e** Distribution of the genes’ average RNA-seq expression for cell cycle genes in G1/S (n = 388), S (n = 835), G2 (n = 183), G2/M (n = 68), and M/G1 (n = 329) phases. Significant differences for each phase group against S phase were determined by Student’s *t*-test (unpaired, two-tailed) assuming equal variances and *p*-values were Bonferroni corrected for multiple testing (**** *p* ≤ 0.0001). **f** RNA-seq profiles for cell cycle genes *CCNB1, CCNE2, PCNA*, and *TOP2A*. **g** Relative expression profiles for *CCNB1, CCNE2, PCNA*, and *TOP2A* as measured by RT-qPCR in two new biological replicates (Val1, Val2; Additional file 1: Fig. S3). Spearman’s correlation coefficients (ρ) were calculated based on the mean RNA-seq expression per time point (Epi1, Epi2) and mean RT-qPCR fold change per time point (Val1, Val2).

Genes that had a cell cycle profile were identified using PLS regression, as described in [1]. We identified 1803 genes with significant periodic expression patterns during the HaCaT cell cycle. These genes are referred to as cell cycle genes. Using the Ensembl BioMart tool (human genes dataset GRCh38.p13), we found 1692 protein coding genes and 108 ncRNAs (Supplementary Dataset 1). Three genes were not detected in BioMart. We used existing annotations [2] to subdivide the genes into five main groups that represent G1/S, S, G2, G2/M, and M/G1 phases of the cell cycle. Of the 1803 cell cycle genes, 835 were assigned to S phase, whereas only 68 were assigned to G2/M phase (Figure 1c); the genes’ cyclic expression patterns are presented in Figure 1d. Examining the RNA-seq expression for the 1803 cell cycle genes by their different cell cycle phases, we observed a significant difference in gene expression between different phases (analysis of variance (ANOVA) *p*-value < 2e-16) and that S phase genes had a significantly lower expression level than the genes expressed in the other phases (Figure 1e). This was in accordance with our previous findings [1]. When examining the genes with comparable expression levels to the cell cycle genes (that is, mean expression ≥ the cell cycle gene with lowest mean expression) but no significant cell cycle-dependent expression pattern (!CC genes), we found that these genes had a bimodal distribution of expression levels (Additional file 1: Fig. S1). We therefore divided this group of genes into genes with high expression level (!CC_high, n = 9259) and genes with low expression level (!CC_low, n = 3003) as reference sets for comparison against the cell cycle genes in further analysis.

To further examine cell cycle profiles at the single gene level, we selected four genes with cell cycle-specific expression patterns and known functions in the cell cycle: Proliferating cell nuclear antigen (*PCNA*), Cyclin E2 (*CCNE2*), DNA topoisomerase 2-alpha (*TOP2A*), and Cyclin B1 (*CCNB1*). PCNA is essential for DNA replication whereas CCNE2 plays a role in the G1/S transition of the cell cycle. Thus, both *PCNA* and *CCNE2* are G1/S-specific genes. TOP2A is involved in processes such as chromosome condensation and chromatid separation whereas CCNB1 is a regulatory protein involved in mitosis. Thus, both *TOP2A* and *CCNB1* are G2/M-specific genes. For all four genes, our RNA-seq data corresponded with their previously reported cell cycle-dependent expression (Figure 1f). Technical validation by RT-qPCR confirmed the RNA-seq expression patterns (Additional file 1: Fig. S2). As a biological validation of the RNA-seq data, we did two new double thymidine block cell cycle synchronization experiments (Val1, Val2) in HaCaT cells (Additional file 1: Fig. S3). Analyses by RT-qPCR showed a high correlation between the original RNA-seq profiles and the new validation experiments (Figure 1g). Thus, both the biological and technical validation indicated that gene expression profiling by total RNA-seq of HaCaT cells could successfully identify cell cycle genes.

### ChIP sequencing maps dynamic transcriptional responses in HaCaT cell cycle

By gene expression profiling we identified a set of genes with cell cycle-dependent expression patterns in HaCaT cells. We wanted to further characterize the dynamic transcriptional response of these genes during the cell cycle. Specifically, we asked whether the genes’ expression changes were accompanied by similar cell cycle-dependent changes in Pol II occupancy at the genes’ transcription start sites (TSSs). Moreover, we asked whether the histone modifications H3K4me3, associated with actively transcribed genes, and H3K27me3, associated with silenced genes, also changed dynamically with gene expression changes. Using ChIP-seq we measured and quantified genome-wide Pol II, H3K4me3, and H3K27me3 occupancy during two independent synchronization experiments (Epi1, Epi2).

First, we performed a sanity check of our sequencing data, by correlating gene expression levels with histone modification patterns and Pol II occupancy in TSS regions for all genes expressed in our RNA-seq data. As Pol II and H3K4me3 both are linked to actively transcribed genes, we expected a positive correlation with gene expression. In contrast, we expected a negative correlation with gene expression for the H3K27me3 modification. Indeed, we found that enhanced Pol II and H3K4me3 signals correlated well with highly expressed genes (Spearman’s correlation coefficient ρ = 0.61 and ρ = 0.64, respectively), whereas there was a negative correlation between gene expression and H3K27me3 signal (ρ = −0.34) (Figure 2a-c). Similarly, dividing the genes into quantiles based on their RNA-seq expression showed that highly expressed genes had the highest Pol II and H3K4me3 signals, whereas genes that were expressed at a low level or not expressed, had low levels of Pol II and H3K4me3 (Additional file 1: Fig. S4). The opposite was observed for H3K27me3, where genes that were not expressed or expressed at a low level had the highest H3K27me3 signal.

**Fig. 2.**
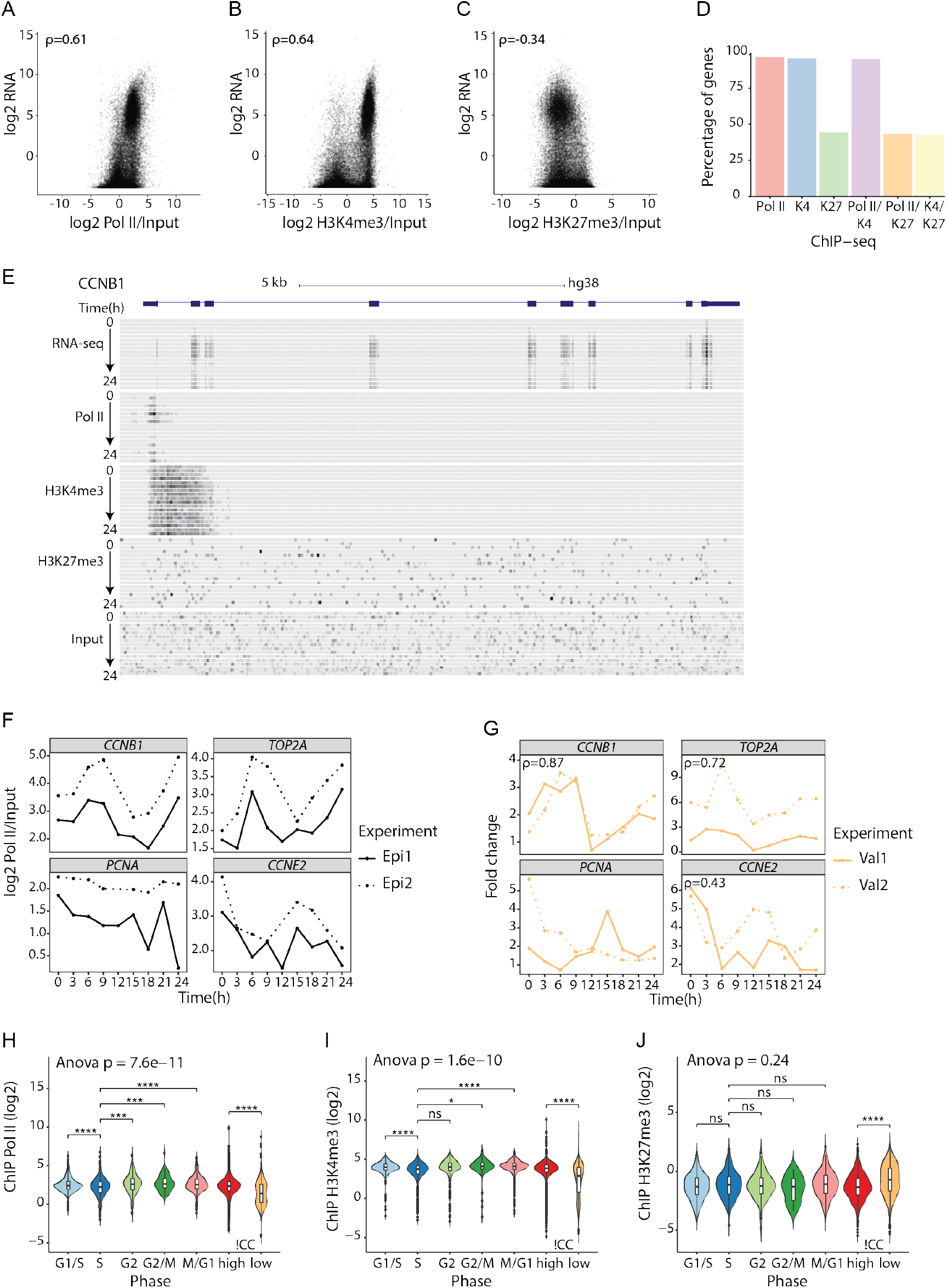
ChIP-seq of synchronized HaCaT cells identifies dynamic H3K4me3, H3K27me3, and PolII changes during cell cycle. **a-c** Average RNA-seq expression (genes expressed>0) plotted against average **(a)** Pol II (n = 30744), **(b)** H3K4m3 (n = 30729), and **(c)** H3K27me3 (n = 30732) ChIP-seq signal normalized against input. Values are Spearman’s correlation coefficients (ρ). **d** Percentage of cell cycle gene promoters containing Pol II, H3K4me3 (K4), or H3K27me3 (K27) marks and combined Pol II/K4, Pol II/K27, or bivalent K4/K27 marks. **e** The UCSC Genome browser (GRCh38/hg38) view of RNA-seq, Pol II, H3K4me3, H3K27me3, and input ChIP-seq data at the *CCNB1* (NM_031966) gene locus. **f** Pol II ChIP-seq profiles for cell cycle genes *CCNB1, CCNE2, PCNA*, and *TOP2A*. **g** Biological validation of Pol II ChIP-seq data for *CCNB1, CCNE2, PCNA*, and *TOP2A*. Spearman’s correlation coefficients (ρ) were calculated based on the mean Pol II ChIP-seq signal per time point (Epi1, Epi2) and mean Pol II ChIP-qPCR fold change per time point from the new biological replicates (Val1, Val2; Additional file 1: Fig. S3). **h-j** Distribution of the genes’ average **(h)** Pol II (n = 1737), **(i)** H3K4me3 (n = 1726), and **(j)** H3K27me3 (n = 802) ChIP-seq signal (normalized against input) for cell cycle genes in G1/S, S, G2, G2/M, and M/G1 phases. Significant differences for each phase group against S phase were determined by Student’s *t*-test (unpaired, two-tailed) assuming equal variances and *p*-values were Bonferroni corrected for multiple testing (ns *p* > 0.05, * *p* ≤ 0.05; ** *p* ≤ 0.01, *** *p* ≤ 0.001, **** *p* ≤ 0.0001).

When investigating the Pol II signal, we observed low signal at time 12 hour in the Epi2 experiment (Additional file 1: Fig. S4), but western blot analysis of the global Pol II protein level throughout the cell cycle indicated no differences in Pol II protein levels between different time points in the cell cycle or between the two independent synchronization experiments (Additional file 1: Fig. S5A). Therefore, we concluded that this was not a biological but a technical issue and decided to exclude the sample (12h, Epi2) in the following analysis. We also investigated global H3K4me3 and H3K27me3 modification levels and found no changes during cell cycle or between the two independent synchronization experiments (Additional file 1: Fig. S5B).

As cell cycle control is a central housekeeping function, we expected the cell cycle genes we identified by RNA-seq to be enriched for H3K4me3 within their promoters, as this epigenetic mark is associated with actively transcribed genes. We also expected the genes to be enriched for Pol II indicating active promoters. Indeed, 96% of the cell cycle genes had promoter regions containing H3K4me3 or Pol II marks. Consistent with these genes being highly expressed, only 45% of the cell cycle gene promoters contained H3K27me3 marks (Figure 2d). In 95% of the gene promoters we found combined Pol II/H3K4me3 marks, whereas 43% of the gene promoters contained combined Pol II/H3K27me3 or bivalent H3K4me3/H3K27me3 marks.

To further inspect the data quality, we visualized RNA-seq and ChIP-seq data in the UCSC Genome browser. *CCNB1* was one of the well-known cell cycle genes we identified, and we observed RNA-seq reads for all exons and Pol II ChIP-seq signals were captured around the TSS (Figure 2e). Additionally, H3K4me3 was highly enriched at this active promoter near TSS, whereas H3K27me3 signals were at the level of the background signal (input ChIP-seq). In contrast, Myelin transcription factor 1 (*MYT1*) is a transcription factor involved in the development of the nervous system, and this gene was not expressed in our HaCaT RNA-seq data (Additional file 1: Fig. S6). The H3K27me3 histone modification was captured at the *MYT1* locus, consistent with this gene not being expressed. Pol II and H3K4me3 signals were observed, but not higher than the level of the background signal.

To characterize the cell cycle ChIP-seq profiles, we first focused on the four cell cycle genes, *CCNB1, CCNE2, PCNA*, and *TOP2A*. The Pol II ChIP-seq profiles for *CCNB1* and *TOP2A* indicated these genes had the highest Pol II signal and active transcription at 9 and 24 hours (Figure 2f), which is consistent with both genes having increased expression in the G2/M phase of the cell cycle (Figure 1f). Additionally, H3K4me3 signals for *CCNB1* and *TOP2A* showed the same cyclic patterns peaking in G2/M phase (Additional file 1: Fig. S7A). *CCNE2* and *PCNA* are both G1/S-specific genes, but only CCNE2 showed the expected Pol II (Figure 2f) and H3K4me3 (Additional file 1: Fig. S7A) profiles peaking at 0-3 and 15-18 hours. As PCNA displayed inconsistent RNA-seq and ChIP-seq profiles, we inspected the sequencing data for the *PCNA* locus in the UCSC Genome browser (Additional file 1: Fig. S8). Interestingly, *PCNA* has two different transcript variants separated by 6,664 bp between their TSSs. As illustrated in Additional file 1: Fig. S8, RNA-seq reads were from the short transcript variant (NM_182649) with a cyclic expression pattern peaking in the G1/S phase. In the ChIP-seq data analysis, the longest transcript variant is always selected (see Methods). Thus, the ChIP-seq signals we observed for *PCNA* were the weak signals from the longest transcript variant (NM_002592).This explains the inconsistency between RNA-seq and ChIP-seq data at the *PCNA* locus, and could also be the case for other genes with multiple TSSs separated by more than 5,000 bp. The H3K27me3 ChIP-seq profiles for *CCNB1, CCNE2, PCNA*, and *TOP2A* showed negative H3K27me3 signal values, which are consistent with these genes being actively transcribed and highly expressed (Additional file 1: Fig. S7B).

To validate the Pol II ChIP-seq data for *CCNB1, CCNE2, PCNA*, and *TOP2A* we did Pol II ChIP-qPCR from the two new cell cycle synchronization experiments (Val1, Val2) in HaCaT cells. We designed PCR primers against the TSS region for these genes (Additional file 1: Table S1); for *PCNA* we designed primers against the short transcript variant (NM_182649). For *CCNB1, TOP2A*, and *CCNE2* we found good correspondence between the original Pol II ChIP-seq experiments and the Pol II ChIP-qPCR from the new validation experiments (Figure 2g). Moreover, the Pol II ChIP-qPCR profile for the short *PCNA* transcript showed increased expression in the G1/S phase (Figure 2g), which is consistent with the RNA-seq (Figure 1f) and RT-qPCR (Figure 1g) expression profiles for *PCNA* peaking in G1/S phase (Additional file 1: Fig. S9).

Since we observed a significant difference in gene expression between the S phase genes and the genes upregulated in the other cell cycle phases (Figure 1e), we wanted to examine the distribution of Pol II, H3K4me3, and H3K27me3 ChIP-seq signals during cell cycle. We included cell cycle genes with positive ChIP-signal in at least two time points of the cell cycle. As for gene expression, we observed a significant difference in both Pol II and H3K4me3 signals between different cell cycle phases, with ANOVA *p*-values of 7.6e-11 and 1.6e-10, respectively. And as for gene expression, the S phase genes had significantly lower Pol II and H3K4me3 signals than the genes expressed in the other phases (Figure 2h-i), with the exception of H3K4me3 signal for genes in G2 phase, which showed no significant difference from S phase genes. As expected, genes with high expression (!CC_high) showed significantly higher levels of Pol II and H3K4me3 signals than genes with low expression (!CC_low). There were no significant differences in H3K27me3 signal between different cell cycle phases, and as expected, !CC_low genes had significantly higher levels of H3K27me3 signal than !CC_high genes (Figure 2j).

### A set of cell cycle genes is highly correlated with Pol II and H3K4me3 changes and has strong enrichment for cell cycle functions

Having quantified gene expression and mapped histone modification patterns together with Pol II occupancy during the cell cycle, we did an integrated bioinformatic analysis of RNA-seq and ChIP-seq data for the cell cycle genes. Specifically, we asked to what extent the genes’ RNA-seq expression through the cell cycle were correlated with their ChIP-seq data and whether such correlation patterns were related to the genes’ functional role in the cell cycle. As a reference, we included the genes without cell cycle-dependent expression profiles, divided by their gene expression levels (!CC_high, !CC_low; Additional file 1: Fig. S1).

We found that all three gene sets (cell cycle (CC) genes, !CC_high, and !CC_low) on average showed a significant positive correlation for RNA-seq expression against Pol II ChIP-seq signal (Figure 3a). Importantly, CC genes showed a significantly higher correlation than !CC genes. The cell cycle genes *CCNB1* (Spearman’s ρ = 0.47), *CCNE2* (ρ = 0.74), and *TOP2A* (ρ = 0.78) are examples of genes with high correlation values. We also found that CC genes and !CC_high genes, but not !CC_low genes, showed a significant positive correlation for RNA-seq expression against H3K4me3 ChIP-seq signal (Figure 3b). Again, CC genes showed a significantly higher correlation than !CC genes. Correlation values for *CCNB1, CCNE2*, and *TOP2A* were ρ = 0.74, ρ = 0.24, and ρ = 0.58, respectively. There were no significant differences between the three gene sets for the H3K27me3 modification (Additional file 1: Fig. S10).

**Fig. 3.**
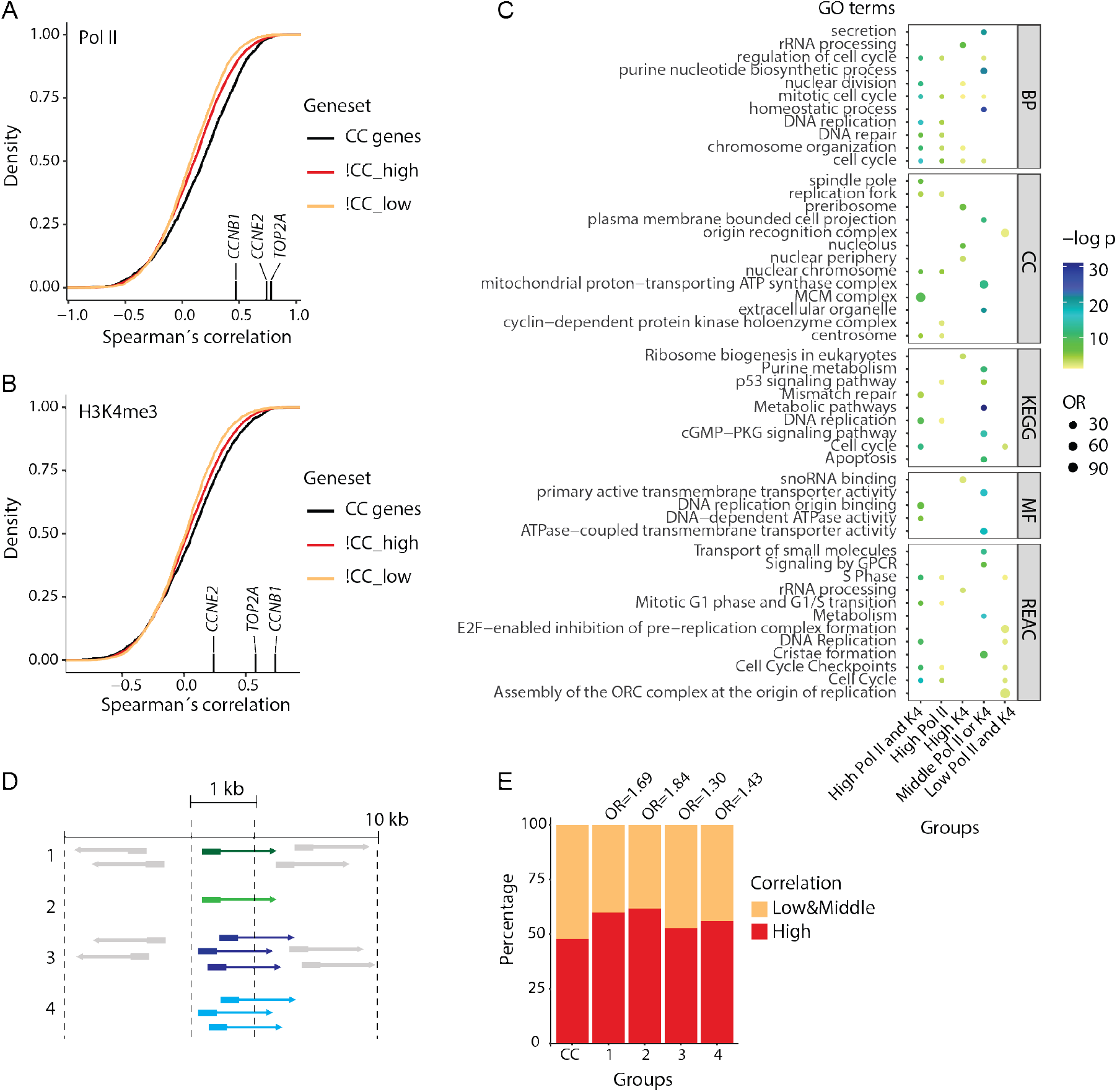
A set of cell cycle genes correlate with PolII and H3K4me3 signals. **a-c** Cumulative distributions of Spearman’s correlation between RNA-seq expression and ChIP-seq signal through the cell cycle for the cell cycle (CC) genes, and other highly (!CC_high) and lowly (!CC_low) expressed genes. Marked on the x axis are the correlation values for *CCNB1, CCNE2*, and *TOP2A*. **(a; Pol II)** All three gene sets had a significant (*p*-value < 2.2e-16) positive correlation for RNA-seq against Pol II ChIP-seq. CC genes showed significantly higher correlation than !CC_high (p = 1.49e-11) and !CC_low (*p*-value < 2.2e-16). **(b; H3K4me3)** CC genes and !CC_high had a significant positive correlation for RNA-seq against H3K4me3 ChIP-seq with *p*-values of 3.83e-15 and < 2.2e-16, respectively. *p*-value for the !CC_low gene set was not significant (p = 0.2287). CC genes showed significantly higher correlation than !CC_high (p = 0.0003859) and !CC_low (p = 1.002e-08). Significant differences were determined by Student’s *t*-test (unpaired, two-tailed) assuming unequal variances. c GO analysis for cell cycle genes. The results show GO biological process (BP), cellular component (CC), molecular functions (MF) terms, and KEGG and REACTOME pathways significantly enriched (*p*-values < 0.05) for cell cycle genes divided in five groups (Additional file 1: Fig. S11). **d** Illustration of the four groups: 1: gene has single TSS (n = 127), 2: gene has single TSS and no other TSSs within 10kb (n = 61), 3: gene has all TSSs within 1 kb (n =356), 4: gene has all TSSs within 1kb and no other TSSs within 10kb (n = 152). **e** Odds ratios from Fisher’s exact tests comparing the fraction of highly correlated genes in the groups (d) with all cell cycle genes with Pol II signals (n = 1735). Group 1 *p* = 0.006, group 2 *p* = 0.026, group 3 *p* = 0.028, and group 4 *p* = 0.041.

Further, we divided the cell cycle genes in high (ρ > 0.2), low (ρ < −0.2), and middle (ρ > −0.2 and ρ <0.2) correlated genes, based on the genes’ Spearman correlation value. We did a gene ontology (GO) analysis for five different groups (Additional file 1: Fig. S11); high correlation for both Pol II and H3K4me3 (n = 400), high correlation for Pol II only (n = 423), high correlation for H3K4me3 only (n = 186), middle correlation for Pol II or H3K4me3 (n = 580), and low correlation for both Pol II and H3K4me3 (n = 125). We identified non-redundant terms among the top 20 significant terms within each GO category. The results showed that genes with high correlation for Pol II and H3K4me3 were specifically enriched for cell cycle-related terms, including cell cycle regulation, DNA replication, and nuclear division (Figure 3c, Supplementary Dataset 2). Moreover, middle correlated genes were specifically enriched for cell signalling, including p53 signalling, whereas genes with low correlation were specifically enriched for initiation of replication. For H3K27me3, genes with high or middle correlation were weakly enriched for cell cycle functions whereas genes with low correlations were weakly enriched for the ribosome biogenesis pathway (Additional file 1: Fig. S12; Supplementary Dataset 3).

The combined RNA-seq and ChIP-seq analysis indicated distinct functions for cell cycle genes based on their correlation with Pol II and H3K4me3 signals, as highly correlated genes had strong enrichment for cell cycle functions. Nevertheless, we wondered to what extent the results were affected by ambiguous mapping of ChIP-seq signals. Specifically, we wondered if the correlation was affected by whether the gene had one or multiple annotated TSSs and by whether the gene was isolated or had annotated TSSs for neighboring genes within its TSS region. Thus, we divided the cell cycle genes into four different groups based on the arrangement of TSSs and other nearby genes (Figure 3d). Genes in group 1 had one single TSS and other genes within 10 kb, whereas genes in group 2 had one single TSS and no other genes within 10 kb. Genes in group 3 had multiple TSSs all within 1 kb and other genes within 10 kb, whereas genes in group 4 had multiple TSSs all within 1 kb and no other genes within 10 kb. Compared with all cell cycle genes (CC), a larger fraction of genes with one TSS (groups 1, 2) were highly correlated with their Pol II signal (Figure 3e); for genes in groups 3 and 4, the fractions were between those of all CC genes and single TSS genes. Moreover, for genes with clearly defined TSS areas and with no overlap with other genes (group 2 compared to 1 and group 4 compared to 3), there was a slightly larger fraction of highly correlated genes. We found no such differences for H3K4me3 and H3K27me3 modifications (Additional file 1: Fig. S13 and S14), possibly because these signals on average cover wider genomic regions than do Pol II.

Thus, whereas ambiguous gene annotations could explain low correlation levels for some genes with cell cycle functions, including *PCNA*, genes with cell cycle-dependent expression and correlated Pol II or H3K4me3 changes were strongly enriched for known cell cycle functions. This result suggested that these highly correlated genes are prime candidates for functional follow-up, so we focused on lncRNAs among these genes.

### Combined RNA-seq and ChIP-seq analysis identifies cell cycle-associated lncRNAs

By combining RNA profiling with ChIP-seq data, we identified 94 cell cycle lncRNAs (Additional file 1: Table S2). Similar to a set of 57 well-described cell cycle genes [2], these lncRNAs had higher expression in proliferating tissues than in non-proliferating tissues (Figure 4a), further supporting their potential role in proliferation. Thus, the combinational use of RNA-seq and ChIP-seq data enabled us to focus on lncRNAs with strong enrichment for cell cycle functions.

**Fig. 4.**
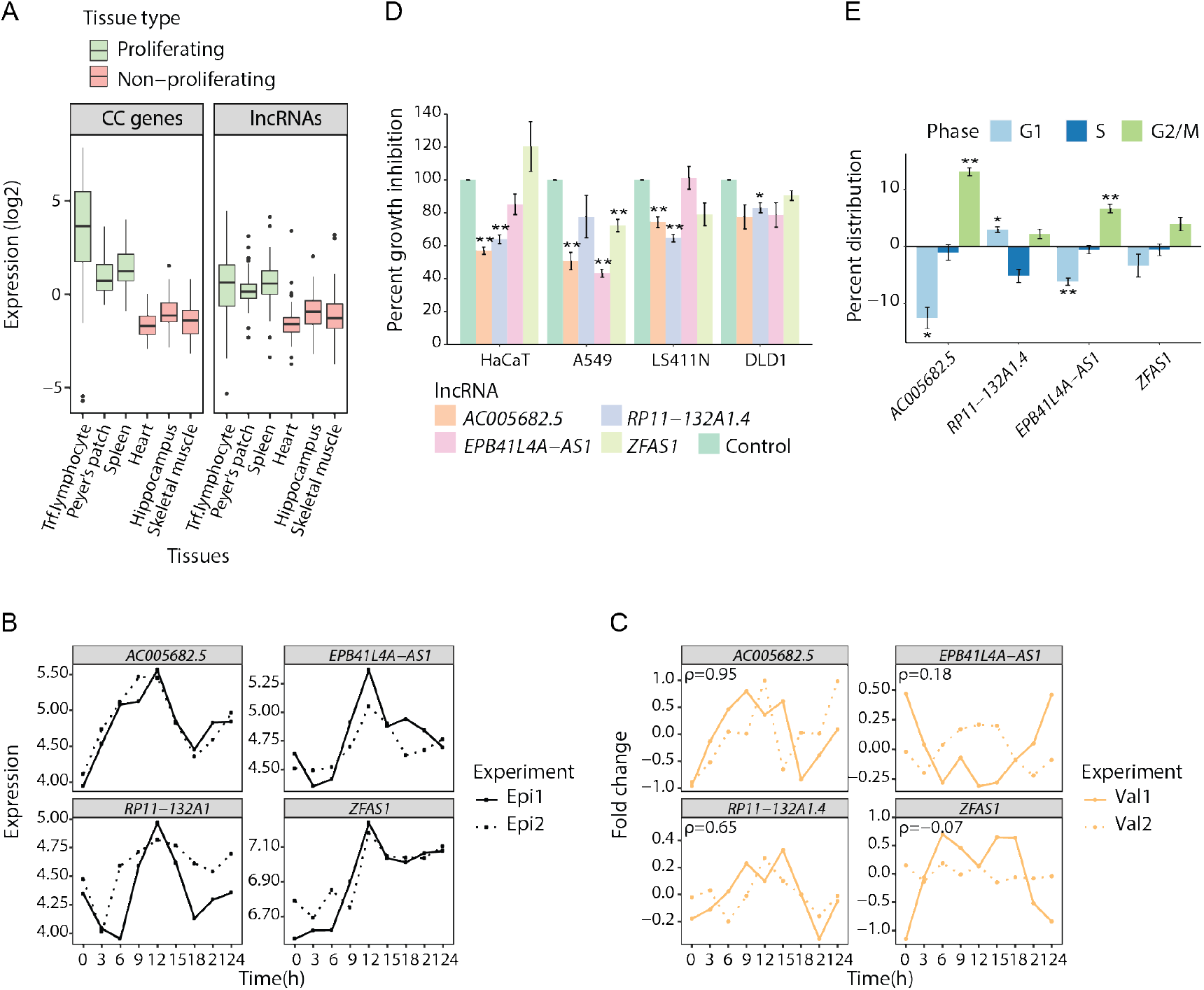
Joint RNA-seq and ChIP-seq analysis identifies lncRNA affecting cell growth and cell cycle progression. **a** Relative tissue expression (log transcript per kilobase million) of known cell cycle genes [2] and our cell cycle lncRNAs in selected tissues from the Genotype-Tissue Expression (GTEx) project. Gene expression values were normalized to relative expression values by subtracting the gene’s average expression across all GTEx tissues. **b** RNA-seq profiles for *AC005682.5, RP11-132A1.4, EPB41L4A-AS1*, and *ZFAS1*. **c** Relative expression profiles for *AC005682.5, RP11-132A1.4, EPB41L4A-AS1*, and *ZFAS1* as measured by RT-qPCR in two new biological replicates. Spearman’s correlation coefficients (ρ) were calculated based on the mean RNA-seq expression per time point (Epi1, Epi2) and mean RT-qPCR fold change per time point (Val1, Val2). **d** Effect of siRNA-mediated knockdown of *AC005682.5* (siRNA A1; siRNA A2 for A549), *RP11-132A1.4* (siRNA R1), *EPB41L4A-AS1* (siRNA E1), and *ZFAS1* (siRNA Z1) on proliferation in four different cell lines. Data are the number of cells following siRNA treatment relative to control-treated cells (percentage of control) as measured by cell counting. Bars and error bars are mean and standard deviation of three or more independent replicates. ANOVA *p*-values from the hierarchical, linear model: *AC005682.5 p* = 7e-4, *RP11-132A1.4 p* = 4e-4, *EPB41L4A-AS1 p* = 0.0896, and *ZFAS1 p* = 0.37. **e** The distribution of cells in G1, S, and G2/M phases in response to knockdown of *AC005682.5* (siRNA A1), *RP11-132A1.4* (siRNA R1), *EPB41L4A-AS1* (siRNA E1), and *ZFAS1* (siRNA Z1) in HaCaT cells. Data are the difference in percentages of G1, S, and G2/M cells of siRNA-treated HaCaT cells to those of control-treated HaCaT cells. Bars and error bars are mean and SD of three independent replicates. **(d-e)** Significant differences were determined by Student’s *t*-test (unpaired, two-tailed) assuming equal variances and *p*-values were Bonferroni corrected for multiple testing (* *p* ≤ 0.05; ** *p* ≤ 0.01).

We selected four lncRNAs for further functional characterization. The two lncRNA candidates *AC005682.5* and *RP11-132A1.4* were chosen as they had high correlation for Pol II and H3K4me3. *AC005682.5* is located between the genes *TOMM7* and *FAM126A* on chromosome 7, while *RP11-132A1.4* is located at the same chromosome between the genes *FIS1* and *IFT22*. The lncRNA *ZFAS1* was identified among the candidates and since it has previously been reported to affect proliferation in different types of cancer [29], we chose this lncRNA as a candidate. *ZFAS1* is located between the genes *DDX27* and *ZNFX1* on chromosome 20. In addition, we chose *EPB41L4A-AS1* (EPB41L4A Antisense RNA 1) as a candidate, as this lncRNA had similar ChIP-seq characteristics as *ZFAS1* (Additional file 1: Fig. S15, Table S2). *EPB41L4A-AS1* is located between the genes *NREP* and *EPB41L4A* on chromosome 5. Tissue-specific expression for the four lncRNA candidates confirmed that all were highly expressed in proliferating tissues (Additional file 1: Fig. S16). All candidates were predicted to have a low protein coding probability, as assessed by the Coding Potential Assessment Tool [30] (Additional file 1: Table S3).

Since lncRNAs are dependent on proximity to their target molecules to exert their function, the subcellular localization can provide important information about their biological role. We used the lncAtlas online tool (https://lncatlas.crg.eu/) to determine the subcellular localization of the four candidate lncRNAs. The lncAtlas displays the subcellular localization of lncRNAs based on a relative concentration index (RCI = concentration of a gene, per unit mass of RNA between cytoplasm and nucleus) derived from RNA-seq data sets from available cell lines and cellular compartments from Human Gencode v24 [31]. Based on RCI from several cell lines, the four lncRNA candidates were mainly localized in either the nucleus or cytoplasm. *AC005682.5* was enriched in the nucleus with an average RCI of −2.03, whereas *RP11-132A1.4, EPB41L4A-AS1*, and *ZFAS1* had the highest level in cytoplasm with an average RCI of 1.82, 1.12, and 1.43, respectively (Additional file 1: Fig. S17).

The RNA-seq data for the lncRNA candidates indicated that *AC005682.5* peaked in G2/M phase, *RP11-132A1.4* and *EPB41L4A-AS1* peaked in M/G1 phase, and ZFAS1 peaked in G1/S phase (Figure 4b). These patterns were confirmed by technical validation by RT-qPCR (Additional file 1: Fig. S18), but only the patterns for *AC005682.5* and *RP11-132A1.4* could be validated in the two new independent double thymidine block synchronization experiments (Val1, Val2) in HaCaT cells (Figure 4c).

To explore the functional role of the candidate lncRNAs in the cell cycle, we used siRNAs to knockdown each candidate (Additional file 1: Fig. S19) and cell counting to evaluate the effect of knockdown on proliferation. We tested four different cell lines – HaCaT, A549, LS411N, and DLD1 – and used a hierarchical, linear model to calculate percent growth inhibition across the four cell lines assuming a random effect for each cell line (Figure 4d). The growth in all four cell lines was significantly affected by *AC005682.5* knockdown, with an average growth inhibition of 35% (Figure 4d). Antisense oligoes are supposedly more effective than siRNAs for nuclear localized transcripts, therefore we used ASO-mediated knockdown to confirm the overall growth inhibitory effect of *AC005682.5* knockdown (average 55%; Additional file 1: Fig. S20). Proliferation was also significantly reduced in all cell lines in response to siRNA-mediated knockdown of *RP11-132A1.4*, with average growth inhibition of 28% and 37% for two independent siRNAs (Figure 4d; Additional file 1: Fig. S20). Meanwhile, the overall growth reduction in all four cell lines in response to *EPB41L4A-AS1* and ZFAS1 knockdown was not significant. However, knockdown of *EPB41L4A-AS1* resulted in a significant growth reduction in HaCaT, A549, and DLD1 cell lines (average 31%; *p*-value = 0.045; Figure 4d), but the growth of LS411N cells was not affected. This trend was validated using an independent siRNA, although the growth reduction in HaCaT, A549, and DLD1 was not significant (*p*-value = 0.070; Additional file 1: Fig. S20). In response to *ZFAS1* knockdown, the growth of the cancerous cell lines A549, LS411N, and DLD1 was significantly reduced by an average of 19% (*p*-value = 0.0054; Figure 4d), which was validated using another siRNA (average 20%; *p*-value = 0.0016; Additional file 1: Fig. S20). Notably, both *ZFAS1* siRNAs gave a slight growth increase in HaCaT cells (20% and 6%; Figure 4d and Additional file 1: Fig. S20).

Finally, we investigated the distribution of HaCaT cells in different cell cycle phases in response to knockdown of the candidate lncRNAs (Figure 4e). Knockdown of *AC005682.5* resulted in a significant reduction of cells in the G1 phase and an enrichment of cells in the G2/M phase (Figure 4e). We validated these overall changes in phase distribution of HaCaT cells using an independent siRNA (Additional file 1: Fig. S21). Knockdown of *RP11-132A1.4* resulted in a significant increased percentage of cells in G1 phase and decreased percentage of cells in S phase (non-significant; ns; Figure 4e). An independent siRNA gave a similar effect on G1 and S phase (Additional file 1: Fig. S21). Knockdown of *EPB41L4A-AS1* resulted in a significant decrease of cells in G1 phase and an enrichment of cells in G2/M phase (Figure 4e), which was validated using an independent siRNA (Additional file 1: Fig. S21). Knockdown of *ZFAS1* resulted in a slight reduction of cells in G1 phase (ns) and an increase of cells in G2/M phase (ns; Figure 4e). This trend was validated using another siRNA (Additional file 1: Fig. S21).

## Discussion

Whereas many lncRNAs have a tissue-specific expression, about 11% of lncRNAs are ubiquitously expressed, suggesting an involvement in cellular functions generally necessary for normal growth and development [32, 33]. By sequencing total RNA through the cell cycle we identified 99 lncRNAs with cell cycle-dependent gene expression. Of these, 48 lncRNAs were highly correlated with changes in Pol II occupancy at their annotated TSS as measured by ChIP-seq, supporting that these lncRNAs are transcribed in a cycle-dependent manner and thereby are likely to have roles in cell proliferation. Indeed, protein coding genes with similar cell cycle-dependent expression and correlated Pol II changes were strongly enriched for cell cycle functions.

We have previously shown that a subset of protein coding cell cycle genes is cell type-specific in their expression and function [1]. Due to the high cell type specificity of many lncRNAs, we therefore expect that some of the lncRNAs identified as cell cycle-associated in HaCaT cells will differ somewhat in other cell types. Nevertheless, similar to protein coding genes with known cell cycle functions, the cell cycle lncRNAs had increased expression in samples from proliferating compared with non-proliferating tissues, suggesting that many of these lncRNAs will have roles in cell proliferation in multiple cell types. Indeed, several lncRNAs that are commonly dysregulated in cancer and that have known cell cycle functions, such as *GAS5, ZFAS1, LINC00963, DANCR*, and *MALAT1* [19, 34–37], were among the 99 lncRNAs with a cyclic expression profile. Moreover, three of the top five lncRNAs (Additional file 1: Table S2) have already been connected to proliferation and cancer in at least one functional study; *CTD-2555C10.3* [38], *SNHG16* [39], and *AC005682.5* [40]. Of these, *SNHG16* is best categorized and is often found overexpressed in cancers where it is associated with poor prognosis [41, 42]. Cao et.al. [39] demonstrated that *SNHG16* increases proliferation in bladder cancer by epigenetic silencing of p21, a potent CDK inhibitor with several functions in the cell cycle [43].

To further investigate the cell cycle-associated lncRNAs, we used siRNAs to down-regulate four candidates – *AC005682.5, RP11-132A1.4, EPB41L4A-AS1*, and *ZFAS1* – and evaluated the resulting effects on cell proliferation in four cell lines (HaCaT, A549, DLD1, and LS411N) and cell cycle progression in HaCaT.

*AC005682.5* was the top cell cycle-associated lncRNA candidate, based on its expression and corresponding correlation to Pol II and H3K4me3 ChIP-seq signals. Downregulation of *AC005682.5* led to growth inhibition in all four cell lines and affected the cell cycle distribution of HaCaT cells by increasing the number of cells in G2/M by more than 10 percentage points. Thus *AC005682.5* seems to be necessary for a normal G2/M progression, which is in line with the gene’s expression profile from RNA-seq, where its expression peaked during the G2/M phase. *AC005682.5* is also known as small nucleolar RNA host gene 26 (*SNHG26*), which belongs to a group of lncRNAs called SNHGs that are often found upregulated in cancers, and that have oncogenic functions connected to proliferation and cell cycle progression. Zimpta et. al. [42] reviewed the role of SNHGs focusing on oncogenic properties and potential clinical applications. Overexpression of SNHGs is often correlated with progression and lower overall survival in several cancers, including lung, stomach, bone, esophagus, liver, brain, and colon. Moreover, in line with our results, the oncogenic properties of SNHGs can be effectively impaired by downregulating their expression using RNA interference [42]. *AC005682.5* has been identified as dysregulated in gene expression datasets from different types of cancer [44–46]. The only study involving any functional characterization of *AC005682.5* was a recent publication identifying *AC005682.5* as a direct transcriptional target of the well-known oncogenic *c-MYC* and a mediator of MYC-driven proliferation in human lymphoid cells [40]. Our study is the first to report that *AC005682.5* is necessary for a proper G2 to M transition during cell cycle, and is important for normal proliferation in HaCaT, A549, DLD1, and LS411N cells.

*RP11–132A1.4* expression peaked in G1 phase, and its knockdown gave growth inhibition in all four cell lines, enrichment of cells in the G1 phase by 3-7 percentage points and a corresponding reduction of cells present in the S phase. These results are in line with a previous functional study of *RP11–132A1.4* in A549 cells [47]. In addition to reporting similar growth inhibition and enrichment of cells in G1 phase, the study revealed that *RP11–132A1.4*, there named E2F1 mRNA stabilizing (*EMS*) lncRNA, is a direct transcriptional target of *c-MYC*. Mechanistically, their results suggest that *EMS* modulates E2F1 stability and promotes G1 to S cell cycle progression through *c-MYC*. *RP11-132A1.4* is upregulated in tissue from colon cancer patients in several datasets, and its expression is associated with poor prognosis [48–50]. In one study *RP11–132A1.4* was differentially expressed between patients with early and advanced stage endometrial carcinoma, where increased expression of *RP11–132A1.4* was associated with disease progression [51]. Previous studies together with our results, indicate an oncogenic role of *RP11-132A1.4*, possibly by interfering with the progression from G1 to the S phase of the cell cycle.

In our study, knockdown of *EPB41L4A-AS1* resulted in an overall growth inhibition of HaCaT, A549, and DLD1 cells, and the distribution of HaCaT cells in the different phases of the cell cycle was also affected with an increase of cells in the G2/M phase by 7 percentage points and a decrease in G1 by 6 percentage points. In the RNA-seq data and RT-qPCR technical validation, *EPB41L4A-AS1* expression peaked in M/G1, but this pattern was only reproduced in one of the two biological validation experiments (Val2, Figure 4c). Several gene expression studies have identified *EPB41L4A-AS1* as both over- and underexpressed in cancer, probably depending on type of tissue and stage of progression [52–54]. Functional studies suggest a central role of *EPB41L4A-AS1* in metabolic reprogramming and as a repressor of the Warburg effect in placental tissue of miscarriage [55] and in cancer cells (cervical, breast, bladder, and liver) [56]. Another functional study investigating the role of *miR-146a* on the proliferation of bone marrow-derived mesenchymal stem (BMSC) cells, reported that *miR-146a* interacts with and inhibits the expression of *EPB41L4A-AS1* and SNHG7. Also, overexpression of *EPB41L4A-AS1* increased proliferation and affected the phase distribution of BMSCs, with a reduction of cells in G1/G0 phase and an increased percentage of cells in the S and G2/M phase [57]. The expression of *EPB41L4A-AS1* is higher in colorectal cancer tissue compared to normal tissue, and in line with our results, knockdown of *EPB41L4A-AS1* decreased proliferation in colorectal cancer cell lines HCT116 and SW620 [58]. Based on previous studies, the expression of *EPB41L4A-AS1* and its biological role seem to vary in different types of cells, tissues, and stage of disease, suggesting a potential role as a biomarker for disease progression and as a therapeutic target. Our study is the first to evaluate how *EPB41L4A-AS1* affects the cell cycle and proliferation in HaCaT, A549, and DLD1 cells. The results suggest an oncogenic role of *EPB41L4A-AS1*, possibly by affecting the progression from G2 to M phase.

The function of *ZFAS1* has been evaluated in several papers, and like many other lncRNAs it varies depending on the type of cell line or tissue being investigated. *ZFAS1* was first described as a regulator of mammary development [59]. Later studies identified it as an oncogene upregulated in several cancers, including lung, colon, ovary, glioma, liver, and gastric cancers, but downregulated in breast cancer [60]. In a study from Fan et. al., overexpression of *ZFAS1* resulted in G1/G0 phase arrest in two breast cancer cell lines [19]. In contrast, we found that *ZFAS1* knockdown had no significant effect on HaCaT proliferation and phase distributions. Instead, our results demonstrated reduced proliferation in response to knockdown in the cancerous cell lines A549, DLD1, and LS411N. Consequently, and in line with previous studies describing the oncogenic characters of *ZFAS1*, our results show that *ZFAS1*’s effects on cell proliferation are cell type-specific and may depend on other oncogenic transformations.

Our analyses suggest that by using a positive correlation between the RNA-seq expression profile and H3K4me3 and Pol II signal as a selection criteria, we identify genes that are actively transcribed and highly enriched for cell cycle functions. Although this is a useful method for detecting cell cycle-associated lncRNAs, cyclic lncRNAs with low correlation to ChIP-seq signal should not be dismissed, as they may also be possible candidates for cell cycle involvement. As exemplified by *PCNA*, genes with several transcripts may have a low correlation value if the wrong TSS was selected for ChIP-seq signal analysis. More sophisticated analyses that use the ChIP-seq data to identify the most likely active TSS per gene may eliminate some of these false negatives.

In summary, our results indicate that all four candidate lncRNAs tested in this study influenced both proliferation and cell cycle progression, although the degree of effect varied between the candidates. The top candidate *AC005682.5*, which has a cyclic expression pattern and the highest overall correlation to Pol II and H3K4me3 ChIP-seq signal, did have a consistent effect on overall growth inhibition and cell cycle phase distribution in response to knockdown. Results from our functional evaluation support that our multi-omics method is well suited for identifying lncRNAs involved in the cell cycle.

## Methods

### Cell culture

All cell lines were obtained from the American Type Culture Collection (ATCC) and cultivated in a humidified incubator at 37°C and 5% CO_2_. The human keratinocyte cell line HaCaT was cultured in Dulbecco’s modified Eagle’s medium (DMEM, Sigma-Aldrich, D6419) supplemented with 10% fetal bovine serum (FBS, Sigma-Aldrich, F7524), 2 mM glutamine (Sigma-Aldrich, G7513), 0.1 mg/ml gentamicin (Gibco, 15710049), and 1.25 μg/ml fungizone (Sigma-Aldrich, A2942). For the lung carcinoma cell line A549 we used DMEM supplemented with 10% FBS and 2 mM glutamine, while the colorectal carcinoma cell line LS411N and the colorectal adenocarcinoma cell line DLD1 were cultivated in RPMI 1640 medium (Gibco, A1049101) supplemented with 10% FBS.

### Cell cycle synchronization

HaCaT cells were seeded in 150-mm culture dishes (2×10^6^ cells each dish) and were arrested in the G1/S transition by double thymidine block. Briefly, cells were treated with 2 mM thymidine for 18 hours, released from the arrest for 10 hours and arrested a second time with 2 mM thymidine for 18 additional hours. After blocking, media was replaced and cells were collected every third hour for 24 hours, covering approximately two cell cycles. Unsynchronized cells were used as a reference sample.

### Cell cycle and fluorescence-activated cell sorting (FACS) analysis

We used FACS analysis to determine the cell cycle phase distribution. HaCaT cells were washed twice with preheated PBS and trypsinated for 8 min before collected using cold PBS supplemented with 3% FBS. Then we centrifuged the cells at 4°C for 5 min. The cell pellet was resuspended in 100 μl cold PBS and fixed by adding 1 ml cold (−20°C) methanol dropwise while vortexing at 1600 rpm and stored at 4°C until DNA measurement. Cells were then washed with cold PBS and incubated with 200 μl of DNase-free RNAse A in PBS (100 μg/ml) for 30 min at 37°C before DNA staining with 200 μl of Propidium Iodide (PI, Sigma; 50 μg/ml) at 37°C for 30 min. Cell cycle analyses were performed by using a BD FACS Canto flow cytometer (BD Biosciences). The excitation maximum of PI is 535 nm and the emission maximum is 617 nm. Here, PI-stained cells were excited with the blue laser (488 nm), and the PI fluorescence was detected in the Phycoerythrin (PE) channel (578 nm). PE channel (578 nm). Quantification of cells in each phase was done with the FlowJo software and the percentage of cells assigned to G1, S, and G2/M phases was calculated.

### Total RNA-seq

Total RNA was isolated using the mirVana miRNA Isolation Kit (ThermoFisher Scientific, AM1560) according to the manufacturer’s protocol. Integrity and stability of RNA samples were assessed by Agilent 2100 Bioanalyzer (Agilent Technologies), whereas the RNA concentration and quality were measured on a NanoDrop ND-1000 UV-Vis Spectrophotometer. RNA-seq libraries were prepared using the Illumina TruSeq Stranded Total RNA with Ribo-Zero™ Human/Mouse/Rat kit, according to the manufacturer’s instructions (Illumina; Additional file 2: Supplementary Method 1). The sequencing (50 cycles single end reads) was performed on an Illumina HiSeq2500 instrument, in accordance with the manufacturer’s instructions. FASTQ files were created with bcl2fastq 2.18 (Illumina).

### Identifying cell cycle genes

RNA-seq raw reads were quality-filtered using fastq_quality_filter 0.0.13 (http://hannonlab.cshl.edu/fastx_toolkit/; parameters −Q33 −q 20 −p 80 −z), and subsequently aligned to human genome (version GRCh38.p7) with STAR 2.4.0.f1 [61]; parameters -- chimSegmentMin 30 --runThreadN 12 --outFilterMultimapNmax 20 --alignSJoverhangMin 8 -- alignSJDBoverhangMin 1 --outFilterMismatchNmax 10 --outFilterMismatchNoverLmax 0.04 -- alignIntronMin 20 --alignIntronMax 1000000). Read alignments were then feature-counted on annotated exons and summarized on genes, using htseq-count 0.6.0 [62]; parameters -r pos -i gene_id -t exon -s yes. The resulting raw count matrix was stripped off genes with zero-count in any of the profiles, preventing such genes from dominating the partial least squares regression (PLS) model. Specifically, out of 58051 gtf-annotated genes in Human Gencode v24, at least a single read was present for 31433 genes, and a total of 14059 was left for analysis. This filtered count matrix was finally transformed to the logarithmic domain and adjusted with precision weights to reduce heteroscedasticity based on abundance using voom from the Limma package [63]. To identify genes with a cell cycle-dependent profile, we used PLS as previously described [1], except that we used the FACS cell fraction matrix directly for the response matrix. Cell cycle phases were assigned as described in [1].

### Quantitative reverse transcription PCR (RT-qPCR)

We isolated total RNA by using the *mir*Vana miRNA Isolation Kit (ThermoFisher Scientific, AM1560) before DNA was removed using TURBO DNA-free™ Kit (Invitrogen, AM1907), according to the manufacturer’s instructions. RNA concentration and quality were measured on a NanoDrop ND-1000 UV-Vis spectrophotometer. Total RNA was reverse transcribed using TaqMan reverse transcription reagents (Applied Biosystems, N8080234) followed by quantitative real-time PCR using SYBR™ Select Master Mix (Applied Biosystems, 44729199) and quantification by the Step One Real-time PCR system (Applied Biosystems). RT^2^ lncRNA PCR assays (Qiagen, 330701) and QuantiTect primer assays (Qiagen, 249900) that were used for lncRNA and mRNA expression analysis are listed in Additional file 1: Table S4. The relative expression of mRNAs and lncRNAs was calculated using the ΔΔCt method [64] with Glyceraldehyde 3-phosphate dehydrogenase (*GAPDH*) as an endogenous control.

### ChIP-seq

Chromatin immunoprecipitation (ChIP) was performed as described in Additional file 2: Supplementary Method 2. The antibodies used for immunoprecipitation (IP) were obtained from Diagenode; anti-H3K4me3 (C1541003-50), anti-H3K27me3 (C15410195), and anti-Pol II (C15200004). As a control for successful IP, qPCR was performed using human positive and negative control qPCR primer sets from Active Motif (Additional file 1: Table S5). Immunoprecipitated material and input chromatin were submitted to the Genomics Core Facility (GCF) at Norwegian University of Science and Technology (NTNU) for library preparation (Additional file 2: Supplementary Method 3) and sequencing. ChIP-seq libraries were prepared using the MicroPlex Library Preparation Kit v4 (Diagenode) and the sequencing (50 cycles single end reads) was performed on an Illumina HiSeq2500 instrument, in accordance with the manufacturer’s instructions (Illumina). FASTQ files were created with bcl2fastq 2.18 (Illumina).

### ChIP-seq data analysis

The ChIP-seq FASTQ files were aligned against the reference human genome hg38 with the hisat2 aligner [65]. Using the -k flag, we only kept 1 primary alignment per read. We also disallowed spliced alignments. We then found the reads per kilobase million (RPKM)-coverage of each alignment file for the genes in hg38 using deepTools bamCoverage [66]. The coverage of each ChIP sample was divided by the average of the coverage of the input sample to provide a normalized expression per gene. For each gene in Human Gencode v24 we found the transcription start sites (TSSs) of the genes and the longest transcript variant was chosen. Then we binned the 10,000 bp area around the TSSs into bins of 50 bp. We found the counts of the reads within each bin by extending each read by 75 (half the fragment size). Then we saw which bin overlapped with the point TSS + 75. Each read was only considered to belong to one bin.

### Pol II ChIP-qPCR

Cells were harvested and Pol II ChIP was performed as described in Additional file 2: Supplementary Method 2, except that cells were sonicated using a Bioruptor Pico (Diagenode) for 14 cycles of 30 sec ON / 30 sec OFF in a volume of 300 μl. No digestion with Micrococcal Nuclease was included. For accurate fragment assessment, the shared chromatin was analyzed on a 2% agarose gel. Fragment size was optimized to be 200-500 bp. As a control for successful IP, qPCR was performed using human positive and negative control qPCR primer sets from Active Motif (Additional file 1: Table S5). PCR primers were designed in the TSS region for selected genes (Additional file 1: Table S1). ChIP DNA was diluted 1:2 in TE buffer and qPCR was performed on immunoprecipitated material and input chromatin. We added 2 μl ChIP DNA and 500 nM of each primer to SYBR Select Master Mix (Applied Biosystems) in technical duplicates. Target values from all qPCR samples were normalized with matched input DNA using the percent input method [100*2^ (Adjusted input - Ct (IP)]. The relative expression of selected genes (Additional file 1: Table S1) was normalized against *GAPDH* and *ACTB* (mean).

### Western blot analysis

We used western blot analysis to validate the Pol II protein level, and H3K4me3 and H3K27me3 modification levels in double thymidine blocked synchronized HaCaT cells (Additional file 2: Supplementary Method 4). Primary antibodies for Pol II (C15200004), H3K4me3 (C15410003), and H3K27me3 (C15410195) were obtained from Diagenode.

### RNA interference

All cells were transfected with 20 nM siRNAs or Antisense LNA GapmeR (Antisense oligo; ASO) using Lipofectamine RNAimax (Invitrogen™, 13778030) when seeded, according to the manufacturer’s protocol. Cells were harvested after 48 and/or 72 hours at about 70% confluence. MISSION^®^ siRNA Universal Negative Control #1 (Sigma, SIC001) and the negative control A Antisense LNA GapmeR (Qiagen, LG00000002) were used as controls for siRNAs and ASO, respectively. The producers and sequences of siRNAs and ASO are listed in Additional file 1: Table S6. All cell culture experiments were performed in three or more independent experiments, and with siRNAs/ASO targeting two different sequences within the same lncRNA.

### Viability assay

We performed cell counting using Moxi z mini automated cell counter (ORFLO Technologies) to investigate how knockdown of the lncRNA candidates affected cell growth. All four cell lines (HaCaT, A549, LS411N, and DLD1) were seeded in triplicates for each condition in a 24-well tray and counted 72 hours after transfection. Each well was washed twice with preheated PBS and trypsinated for 5-10 min before the cells were resuspended in preheated growth medium and counted. We applied a two-tailed, paired Student’s *t*-test to test whether the growth was significantly different (*p* < 0.05) between cells transfected with negative control siRNA/ASO and a lncRNA target-specific siRNA/ASO in at least three independent experiments.

### Data analysis

All data analyses were performed in R. Plots were made using the packages gglot2 and ggpubr. Gene ontology (GO) analyses were done with the package gProfileR.

## Supporting information

Cell cycle genes

GO analysis H3K27me3

GO analysis Pol II and H3K4me3

Graphical abstract

## Declarations

### Ethics approval and consent to participate

Not applicable

### Consent for publication

Not applicable

### Availability of data and materials

The RNA-seq and ChIP-seq datasets generated and analyzed during the current study are available in the European Nucleotide Archive (https://www.ebi.ac.uk/ena/) under the accession number of PRJEB40813.

### Authors’ contributions

SAH designed and performed the cell cycle synchronization experiments, isolated RNA and performed ChIP, did the bioinformatic and statistical analysis and was a major contributor in writing the manuscript. HS designed and performed wet lab experiments (cell cycle synchronization validation, RT-qPCR, western blot analysis, RNA interference, cell cycle and viability assays), contributed with statistical analysis and was a major contributor in writing the manuscript. AK analyzed RNA-seq data and identified cell cycle genes. EBS analyzed ChIP-seq data. KGN performed cell cycle synchronization validation experiments and performed Pol II ChIP-qPCR analysis. NBL performed all FACS analysis. LCO and KC contributed with bioinformatics and statistical analysis. PAA contributed with cell cycle synchronization experiments. PS designed and supervised the study and edited the manuscript. All authors read and approved the final manuscript.

## Acknowledgements

This work was supported by the Norwegian Cancer Society and The Research Council of Norway. The library preparation and sequencing were provided in close collaboration with the Genomics Core Facility (GCF), Norwegian University of Science and Technology (NTNU). GCF is funded by the Faculty of Medicine and Health Sciences at NTNU and Central Norway Regional Health Authority.

## Authors’ information

Siv Anita Hegre and Helle Samdal contributed equally to this work.

## Supplementary information

**Additional file 1**: Supplementary Figures and Tables

**Additional file 2**: Supplementary Methods

**Supplementary dataset 1**: Cell cycle genes identified in this study

**Supplementary dataset 2**: GO analysis Pol II and H3K4me3

**Supplementary dataset 3**: GO analysis H3K27me3

