## Supplementary Figures and Tables for "Joint changes in RNA, RNA polymerase II, and promoter activity through the cell cycle identify non-coding RNAs involved in proliferation"

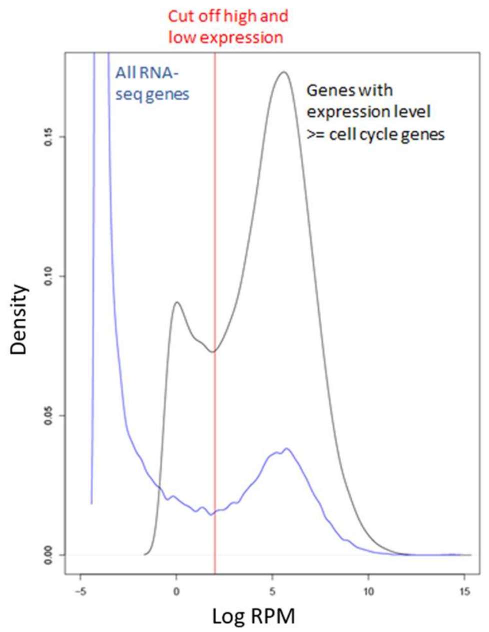


**Supplementary Figure S1: Density plot of the bimodal distribution of RNA-seq expression.** The blue line represents the mean expression for all genes in the RNA-seq data set (n = 65988). The black line represents genes with mean expression ≥ the cell cycle gene with lowest mean expression (n = 14065). This set of genes included cell cycle genes (n = 1803) and genes with no significant cell cycle-dependent expression pattern (!CC genes, n = 12262). The set of genes had a bimodal distribution of expression levels, where the red line represents the cut off between high and low RNA-seq expression. We divided the !CC genes into genes with high expression level (!CC_high, n = 9259) and genes with low expression level (!CC_low, n = 3003).

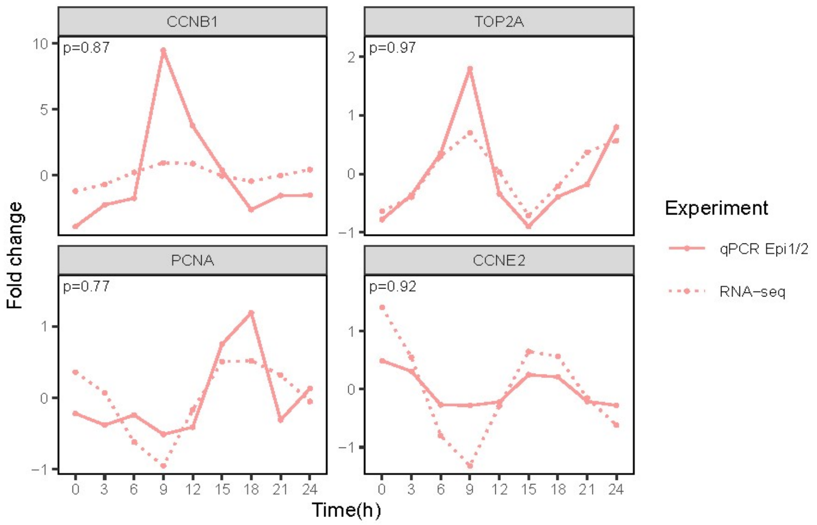


**Supplementary Figure S2: Technical validation of RNA-seq expression for *CCNB1*, *CCNE2*, *PCNA*, and *TOP2A*.** Spearman's correlation coefficients (ρ) were calculated based on the mean RNA-seq expression per time point and mean RT-qPCR fold change per time point from the original experiments (Epi1, Epi2).

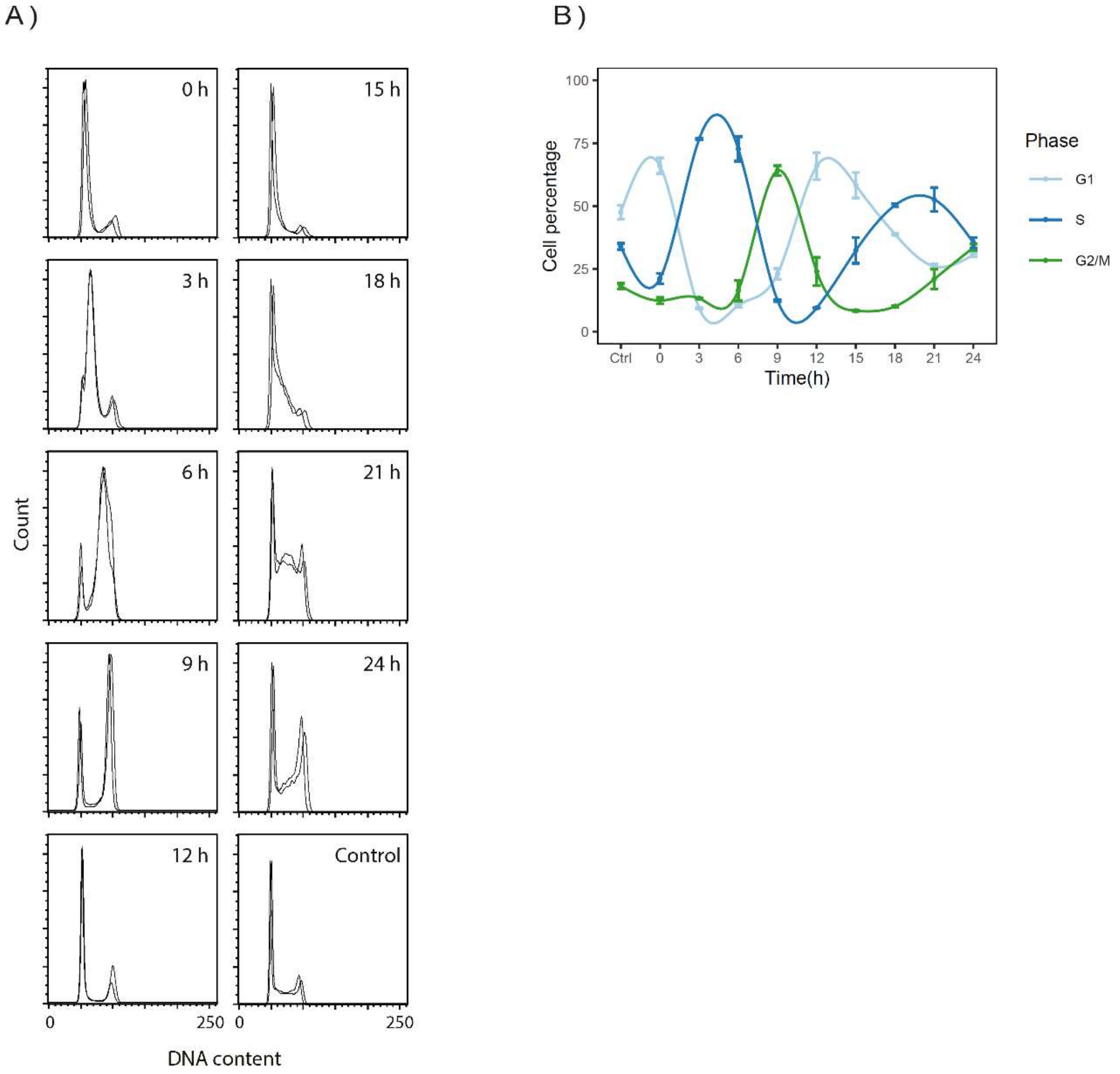


**Supplementary Figure S3: Synchrony of double thymidine blocked HaCaT cells from two validation experiments (Val1, Val2). (A)** Cell synchrony was monitored by flow cytometry of propidium iodide-stained cells. The figure shows superimposed DNA content profiles of the two replicate experiments for each time point. Horizontal axes show DNA content (arbitrary units) and vertical axes show the number of cells with the corresponding DNA content. Control is unsynchronized cells. **(B)** Percentage of cells assigned to G1, S, and G2/M phases for each of the time points analyzed. Values and error bars are averages and standard deviations (n = 2).

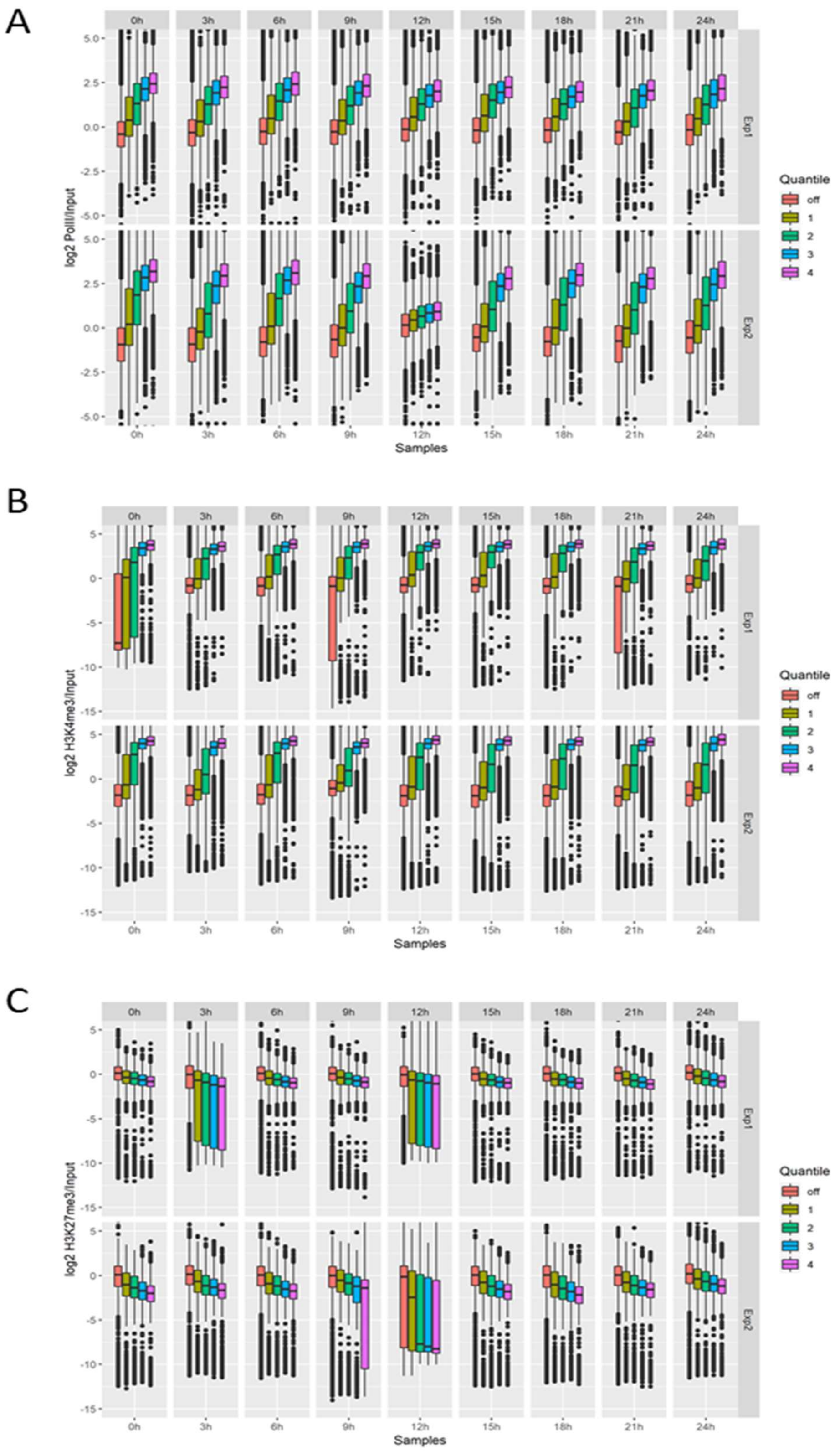


**Supplementary Figure S4: Average abundance of ChIP-seq signal in promoter regions around the transcription start site (TSS).** The sum of counts within each bin of size 50 around the TSS area of all annotated genes. We divided the genes into quantiles based on RNA-seq expression; off = genes not expressed, 1 = 0-25%, 2 = 25-50%, 3 = 50-75%, 4 = 75-100%. **(A)** Pol II ChIP-seq, **(B)** H3K4me3 ChIP-seq, and **(C)** H3K27me3 ChIP-seq.

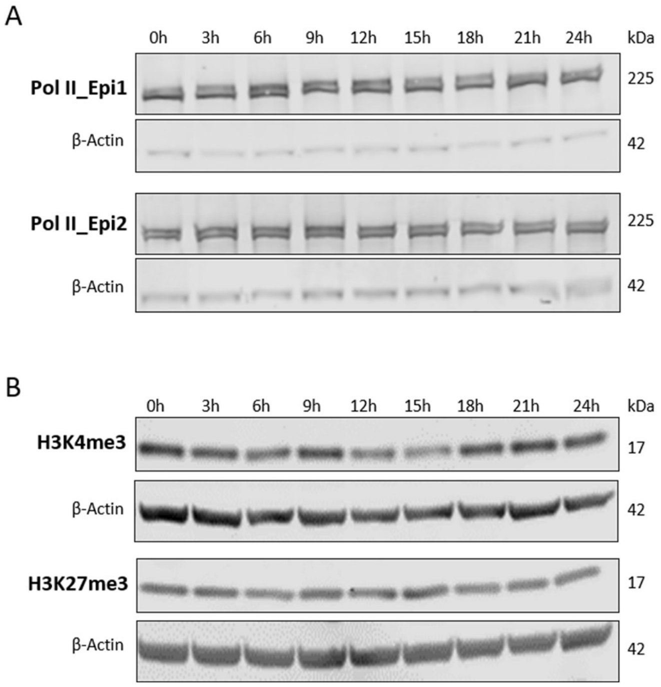


**Supplementary Figure S5: Western blot analysis. (A) Pol II:** Protein was isolated from the original experiments (Epi1,Epi2) for Pol II (225 kDa) detection. **(B) H3K4me3 and H3K27me3:** Protein was isolated from the two validation experiments (Val1, Val2) for H3K4me3 (17 kDa) and H3K27me3 (17 kDa) detection. Quantification (data not shown) indicated no changes during cell cycle or between the different synchronization experiments for Pol II, H3K4me3, or H3K27me3.

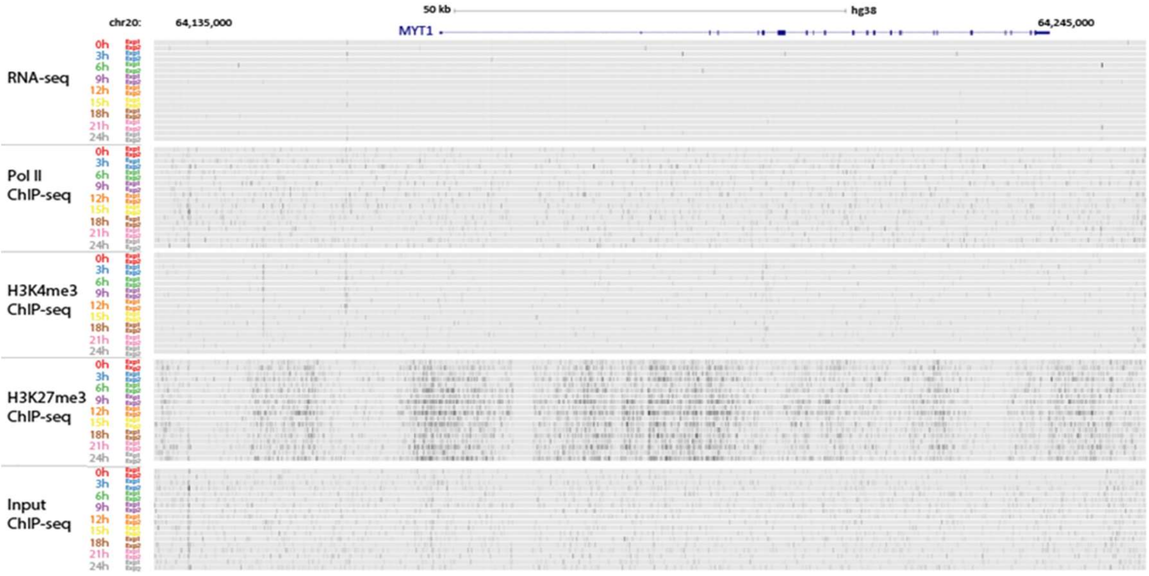


**Supplementary Figure S6: The UCSC Genome browser (GRCh38/hg38) view of RNA-seq and Pol II, H3K4me3, H3K27me3, and Input ChIP-seq data at the *MYT1* (NM_004535) gene locus.**

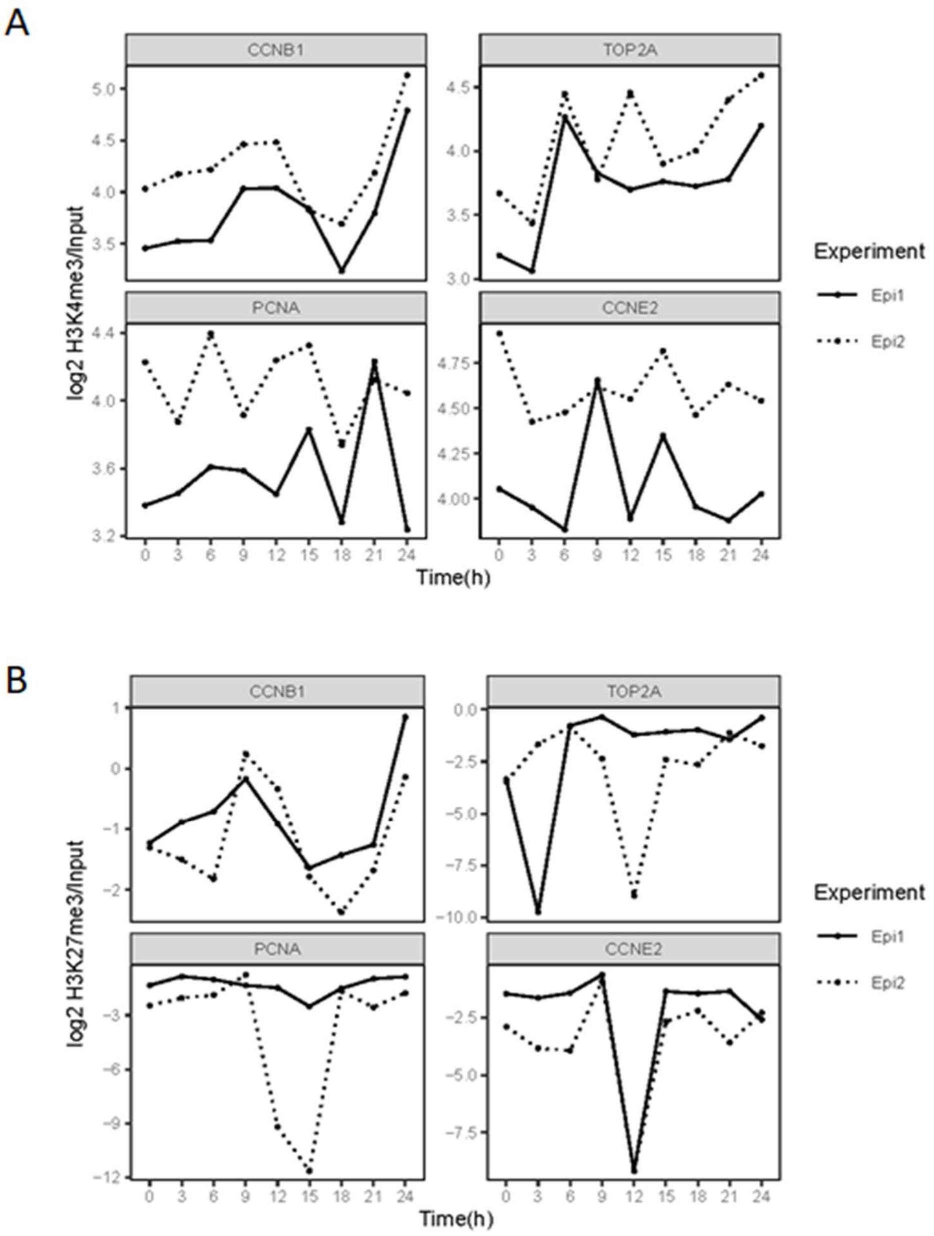


**Supplementary Figure S7: (A) H3K4me3 and (B) H3K27me3 ChIP-seq profiles for the cell cycle genes *CCNB1*, *CCNE2*, *PCNA*, and *TOP2A*.** Negative values on the y axis for H3K27me3 ChIP-seq indicate the ChIP-seq signal is lower than the background (input) signal.

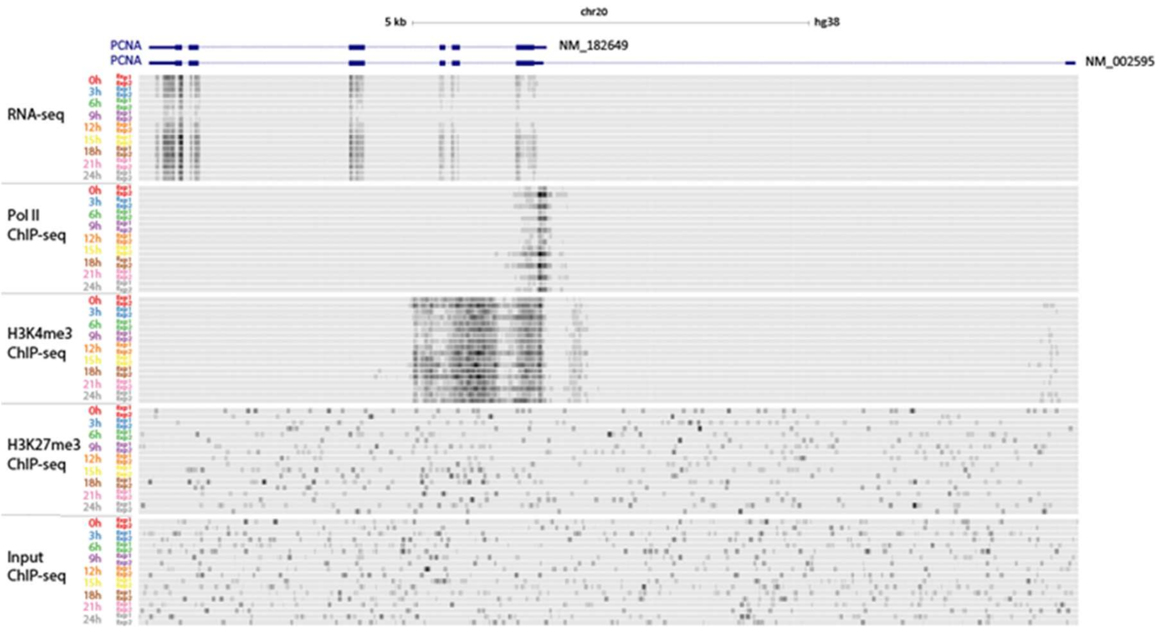


**Supplementary Figure S8: The UCSC Genome browser (GRCh38/hg38) view of RNA-seq and Pol II, H3K4me3, H3K27me3, and Input ChIP-seq data at the *PCNA* (NM_182649 and NM_002592) gene locus.** The Pol II and H3K4me3 data indicate that the short isoform (NM_182649) and not the long isoform (NM_002592) is being transcribed.

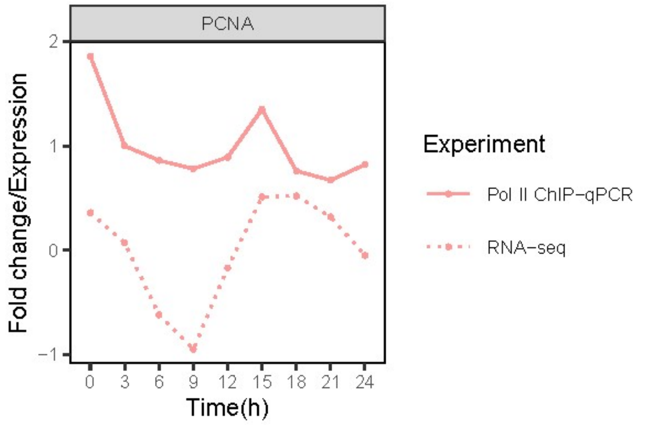


**Supplementary Figure S9: *PCNA* Pol II ChIP-qPCR and RNA-seq expression**. Expression is the mean RNA-seq expression per time point (Epi1, Epi2) and mean Pol II ChIP-qPCR fold change per time point from the new biological replicates (Val1, Val2).

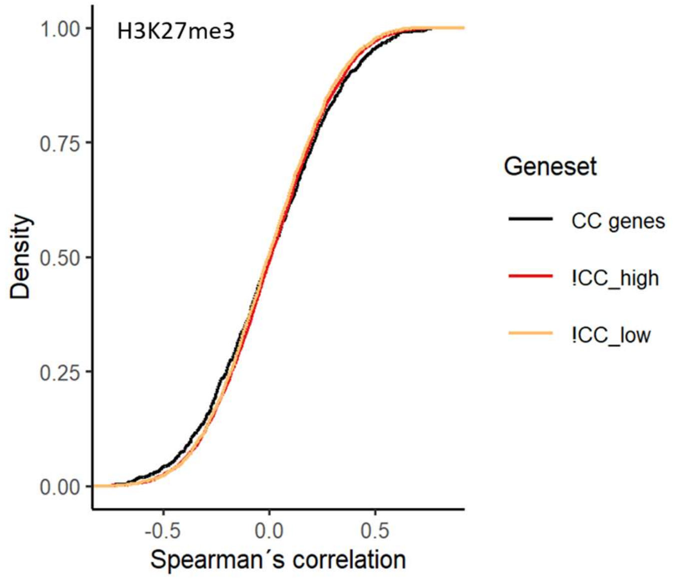


**Supplementary Figure S10: Spearman's correlation analysis of RNA-seq expression against H3K27me3 ChIP-seq signal.** There was no clear shift in the mean correlations for three gene sets CC genes, !CC_high, and !CC_low (*p*-values of 0.63, 0.023, and 0.84, respectively.) There were no significant differences between the gene sets (CC genes against !CC_high *p*-value = 0.72 and CC genes against !CC_low *p*-value = 0.60). Significant differences were determined by Student's *t*-test (unpaired, two-tailed) assuming unequal variances.

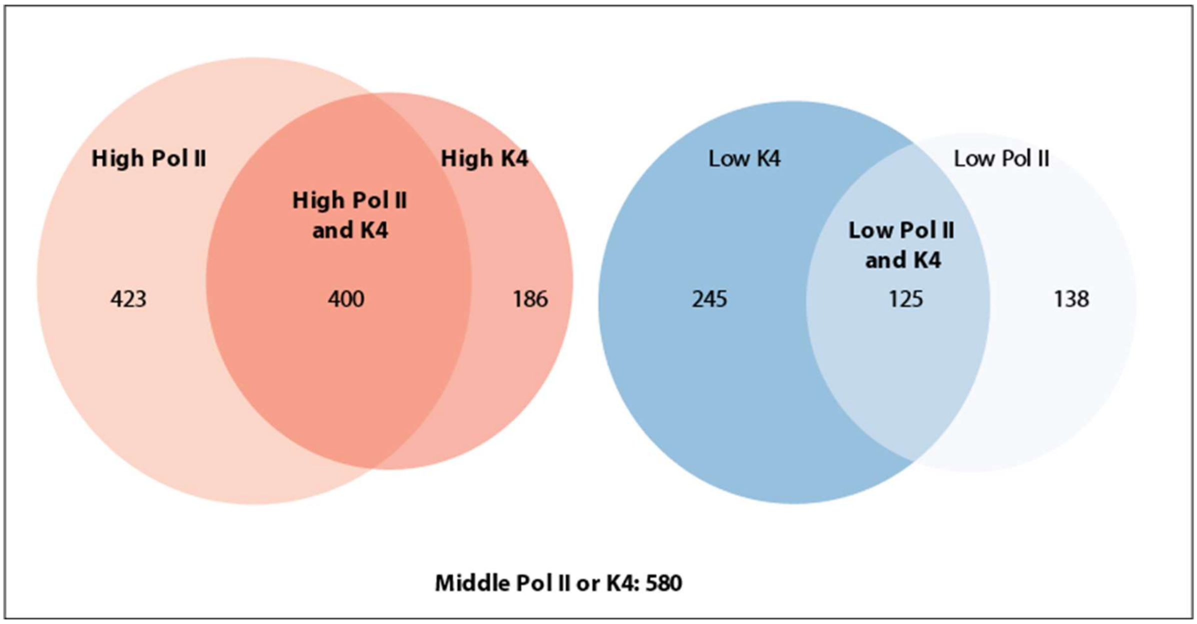


**Supplementary Figure S11: Venn diagram for cell cycle genes with combined Pol II and H3K4me3 (K4) signal (n = 1714).** Genes were divided in high ($\rho > 0.2$), low ($\rho < -0.2$), and middle ($\rho > -0.2$ and $\rho < 0.2$) correlated genes, based on the genes´ Spearman's correlation value for RNA-seq expression against ChIP-seq signal. We did GO analysis for five different groups (in bold); high correlation for Pol II and K4 (n = 400), high correlation for Pol II (n = 423), high correlation for K4 (n = 186), middle correlation for Pol II or K4 (n = 580), and low correlation for Pol II and K4 (n = 125).

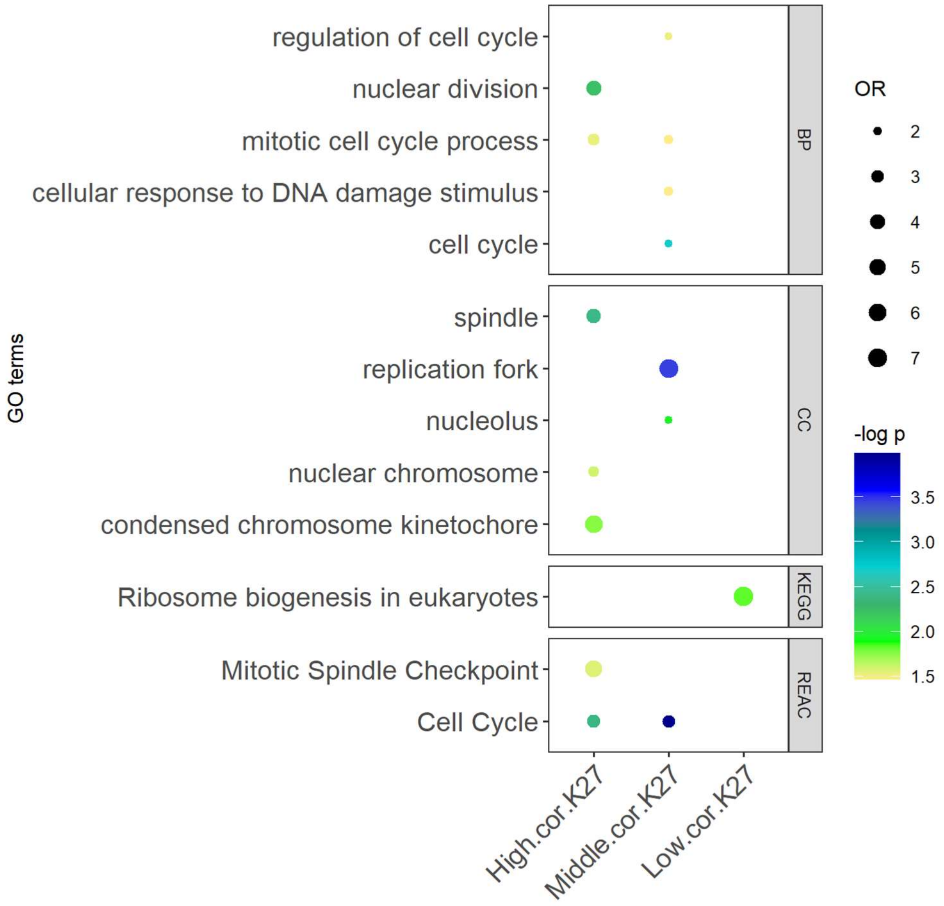


**Supplementary Figure S12: Gene ontology (GO) analysis for cell cycle genes correlated with H3K27me3.** The results show GO biological process (BP), cellular component (CC) terms, and KEGG and REACTOME pathways significantly enriched (*p*-values < 0.05) for cell cycle genes divided in three different groups; high correlation ($\rho > 0.2$), middle correlation ($\rho > -0.2$ and $\rho < 0.2$), and low correlation ($\rho < -0.2$). Correlation values are the results of Spearman's correlation analysis of RNA-seq expression against H3K27me3 signal.

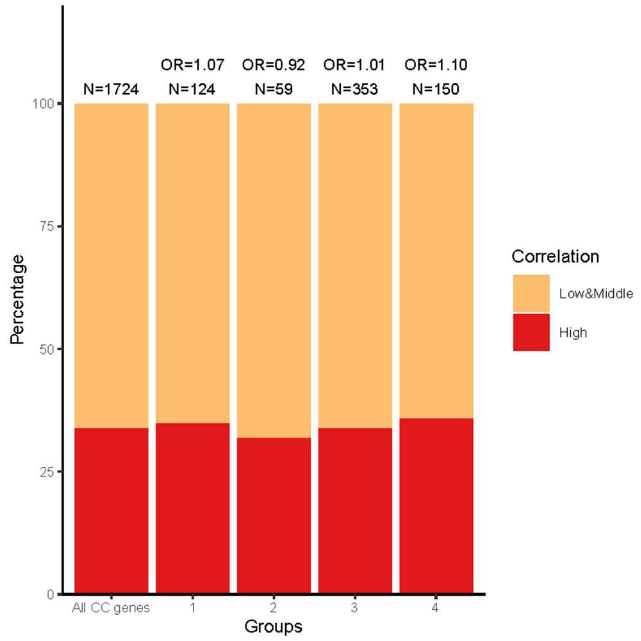


**Supplementary Figure S13: Odds ratios from Fisher's exact tests for H3K4me3**. We compared the fraction of highly correlated genes in the four gene groups (Figure 3D) with all cell cycle (CC) genes with H3K4me3 signals in their TSS (n = 1734). *P*-values (ns): group 1 $p$ = 0.77, group 2 $p$ = 0.89, group 3 $p$ = 0.95, and group 4 $p$ = 0.59.

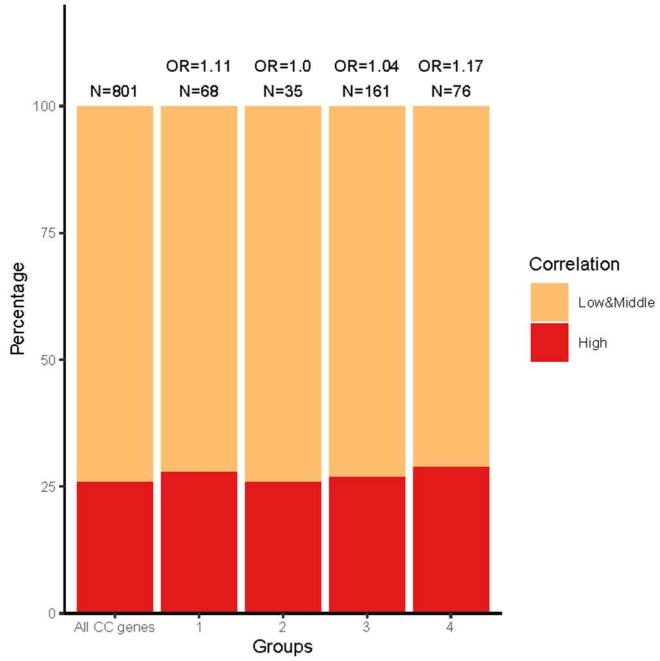


**Supplementary Figure S14: Odds ratios from Fisher's exact tests for H3K27me3**. We compared the fraction of highly correlated genes in the four gene groups (Figure 3D) with all cell cycle (CC) genes with H3K27me3 signals in their TSS (n = 801). *P*-values (ns): group 1 $p$ = 0.77, group 2 $p$ = 1, group 3 $p$ = 0.84, and group 4 $p$ = 0.58.

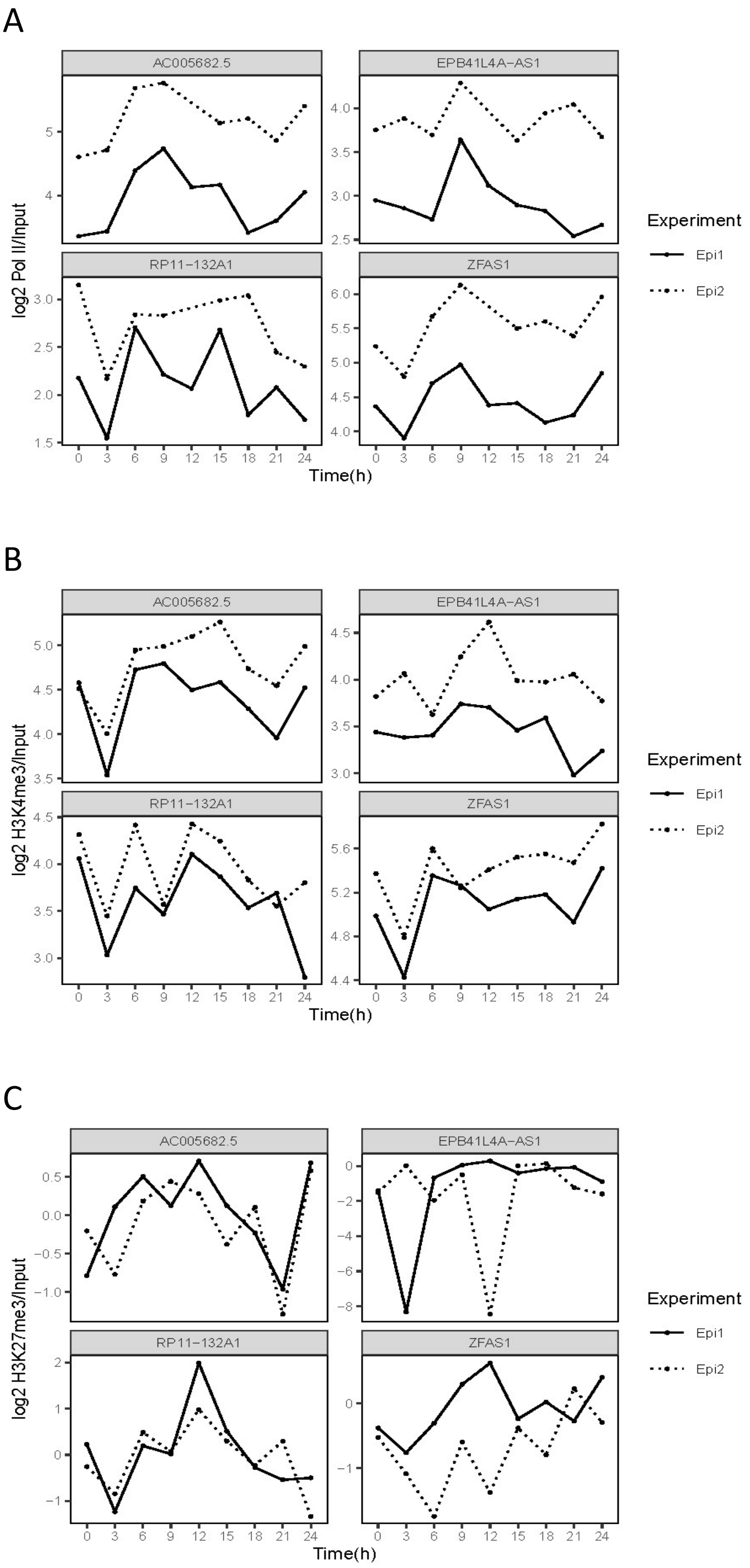


**Supplementary Figure S15: ChIP-seq data for the four lncRNA candidates. (A)** Pol II, **(B)** H3K4me3, and **(C)** H3K27me3 ChIP-seq profiles for *AC005682.5*, *EPB41L4A-AS1*, *RP11-132A1.4*, and *ZFAS1*.

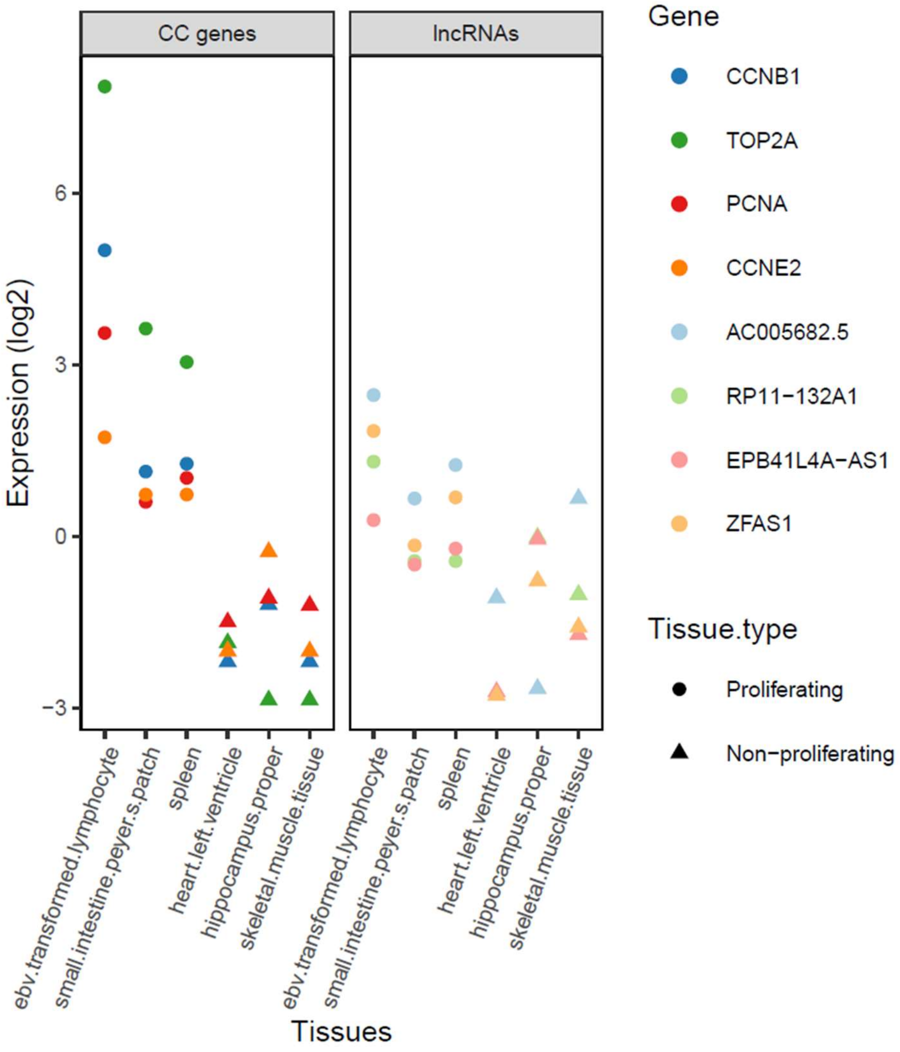


**Supplementary Figure S16: Expression data for known cell cycle genes and the four lncRNA candidates in proliferating and non-proliferating tissues.** Relative tissue expression (log transcript per kilobase million) of known cell cycle genes (1) and our cell cycle lncRNAs in selected tissues from the Genotype-Tissue Expression (GTEx) project. Gene expression values were normalized to relative expression values by subtracting the gene's average expression across all GTEx tissues.

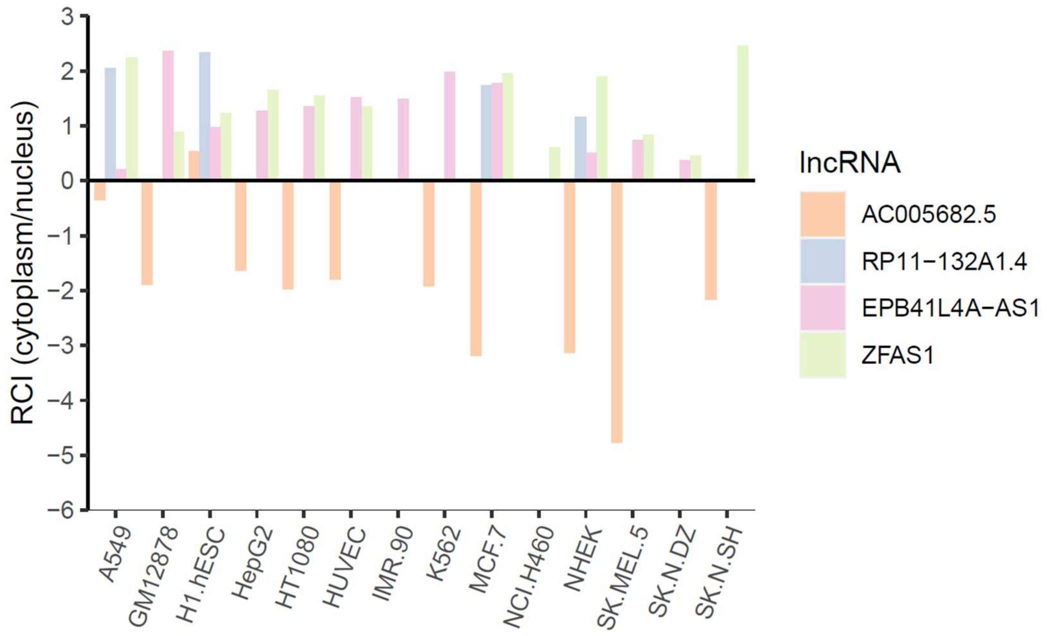


**Supplementary Figure S17: Subcellular localization of lncRNA candidates**. Data are from the lncAtlas (https://lncatlas.crg.eu/) and display the subcellular localization based on a relative concentration index (RCI) of RNA between cytoplasm and nucleus.

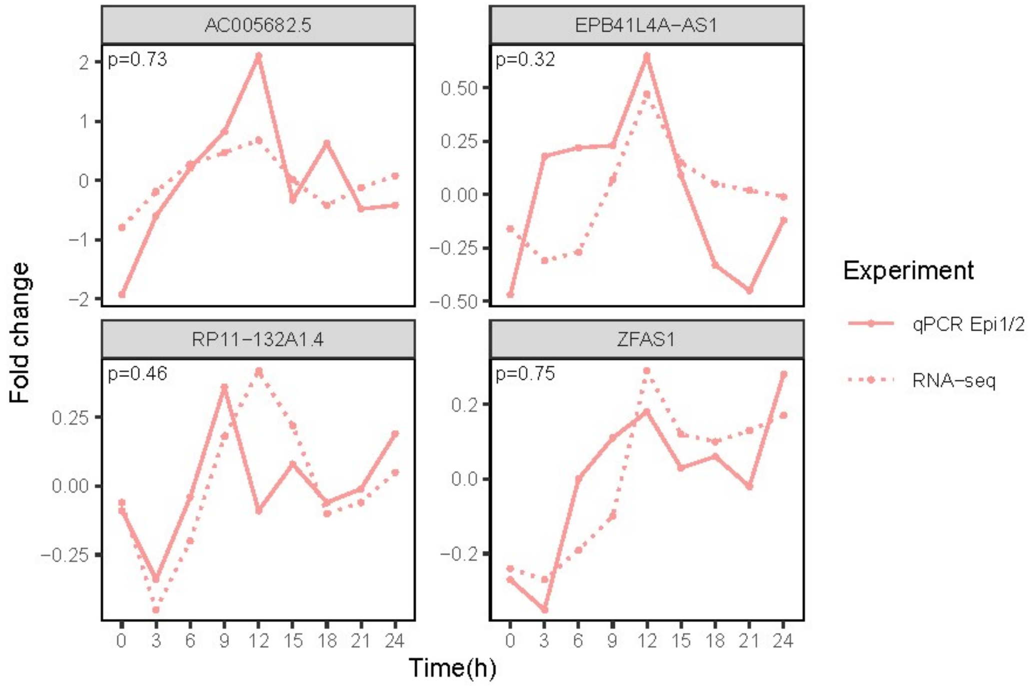


**Supplementary Figure S18: Technical validation of RNA-seq expression for *AC005682.5*, *EPB41L4A-AS1*, *RP11-132A1.4*, and *ZFAS1***. Spearman's correlation coefficients (ρ) were calculated based on the mean RNA-seq expression per time point and mean RT-qPCR fold change per time point from original experiments (Epi1, Epi2).

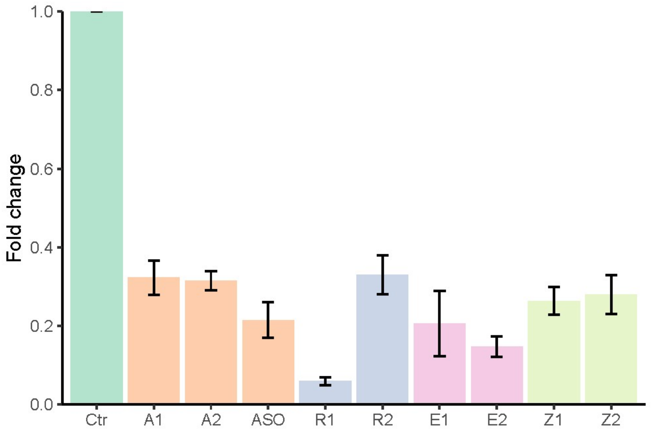


**Supplementary Figure S19: Percentage downregulation of the lncRNA candidates** *AC005682.5* (siRNAs; A1 and A2, and ASO), *RP11-132A1.4* (siRNAs; R1 and R2), *EPB41L4A-AS1* (siRNAs; E1 and E2), and *ZFAS1* (siRNAs; Z1 and Z2) by using two siRNAs with different target sequences. Data are presented as fold change expressions of target lncRNAs following siRNA treatment relative to control-treated cells as measured by RT-qPCR in HaCaT cells. Bars and error bars are mean and standard deviation of two or more independent replicates.

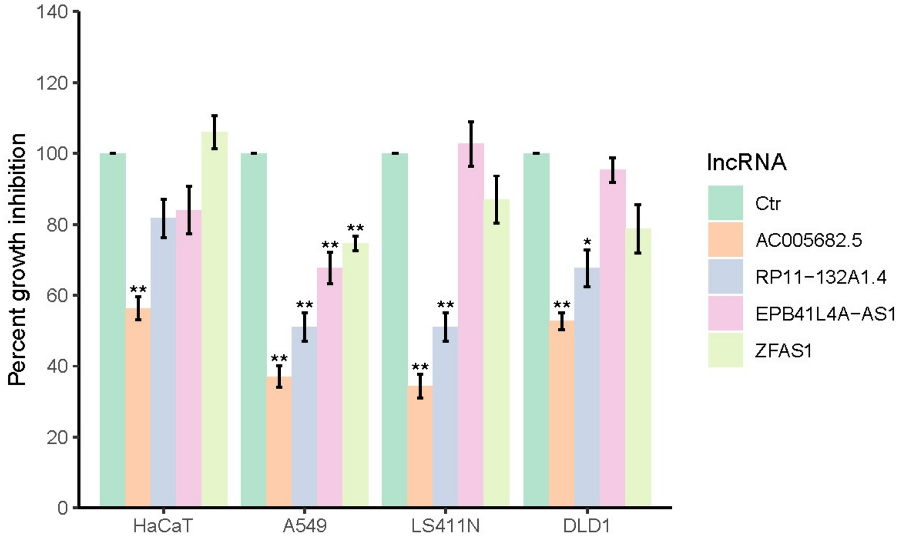


**Supplementary Figure S20: Proliferation assay results.** Effect of siRNA/ASO-mediated knockdown of *AC005682.5* (ASO), *RP11-132A1.4* (siRNA R2), *EPB41L4A-AS1* (siRNA E2), and *ZFAS1* (siRNA Z2) on proliferation in four different cell lines. Data are the number of cells following siRNA treatment relative to control-treated cells (percentage of control) as measured by cell counting. Bars and error bars are mean and standard deviation of three or more independent replicates. Significant differences were determined by Student's *t*-test (unpaired, two-tailed) assuming equal variances and *p*-values were Bonferroni corrected for multiple testing (* $p \leq 0.05$; ** $p \leq 0.01$). ANOVA *p*-values from the hierarchical, linear model: *AC005682.5* $p = 0$, *RP11-132A1.4* $p = 0.001$, *EPB41L4A-AS1* $p = 0.1373$, and *ZFAS1* $p = 0.0902$.

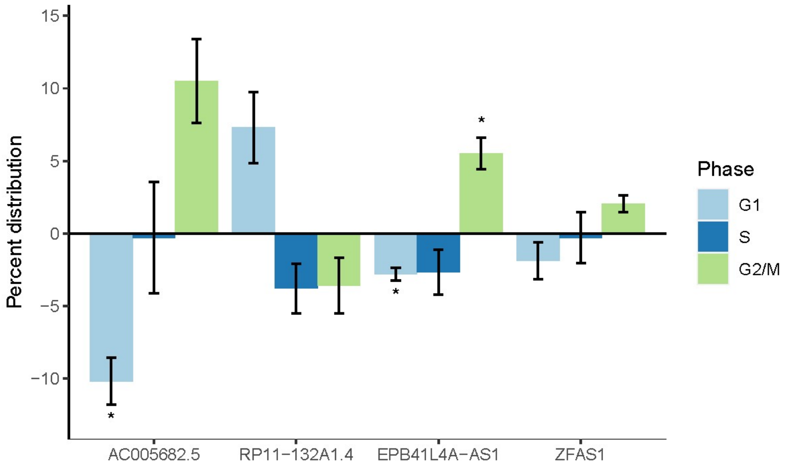


**Supplementary Figure S21: Cell cycle phase distribution results.** The distribution of cells in G1, S, and G2/M cell cycle phases in response to siRNA-mediated knockdown of *AC005682.5* (siRNA A2), *RP11-132A1.4* (siRNA R2), *EPB41L4A-AS1* (siRNA E2), and *ZFAS1* (siRNA Z2) in HaCaT cells. Data are the difference in percentages of G1, S, and G2/M cells of siRNA-treated HaCaT cells to those of control-treated HaCaT cells. Bars and error bars are mean and standard deviation of three independent replicates. Significant differences were determined by Student's *t*-test (unpaired, two-tailed) assuming equal variances and *p*-values were Bonferroni corrected for multiple testing (* $p \leq 0.05$).

### Supplementary tables

**Supplementary Table S1: Primer sequences of selected genes for Pol II ChIP-qPCR validations studies.** S = sense, AS = antisense

| Gene | Sequence | PCR product (bp) |
|---|---|---|
| PCNA | S-5'-TAGCTGGTTTCGGCTTCAGG-3' | 106 |
| | AS-5'-TAAACGGTTGCAGGCGTAG-3' | |
| TOP2A | S-5'-AAGCGACTAAACAGGCAGG-3' | 103 |
| | AS-5'-GGCTAAAGGAAGGTTCAAGTG-3' | |
| CCNB1 | S-5'-ACGAACAGGCCAATAAGGAG-3' | 103 |
| | AS-5'-ACCCAGCAGAAACCAACAG-3' | |
| CCNE2 | S-5'-GCTTTCTTTCCTCCCACATC-3' | 102 |

**Supplementary Table S2: Long non-coding RNAs with RNA-seq/ChIP-seq Spearman's correlation values.** Gene type is from the Human Genes GRCh38.p13 database.

| EnsemblGene | EntrezGene | Phase | Gene type | Cor Pol II | Cor H3K4me3 | Cor H3K27me3 |
|---|---|---|---|---|---|---|
| ENSG00000228649 | AC005682.5 | G2/M | lncRNA | 0.84 | 0.49 | 0.70 |
| ENSG00000255121 | RP11-110I1.12 | S | lncRNA | 0.78 | 0.44 | 0.09 |
| ENSG00000163597 | SNHG16 | M/G1 | lncRNA | 0.76 | 0.68 | NA |
| ENSG00000259230 | CTD-2555C10.3 | S | lncRNA | 0.68 | 0.64 | NA |
| ENSG00000173209 | AHSA2 | S | pseudogene | 0.67 | 0.26 | NA |
| ENSG00000230133 | FLJ33581 | G2 | lncRNA | 0.66 | NA | 0.27 |
| ENSG00000275481 | RP11-474P2.6 | S | lncRNA | 0.66 | 0.55 | NA |
| ENSG00000251669 | FAM86EP | G1/S | pseudogene | 0.64 | 0.46 | 0.10 |
| ENSG00000257167 | TMPO-AS1 | S | lncRNA | 0.64 | -0.03 | NA |
| ENSG00000182796 | TMEM198B | S | pseudogene | 0.62 | 0.13 | 0.12 |
| ENSG00000263753 | LINC00667 | S | lncRNA | 0.58 | 0.35 | 0.62 |
| ENSG00000256268 | RP11-221N13.3 | S | lncRNA | 0.55 | -0.26 | NA |
| ENSG00000238105 | GOLGA2P5 | S | pseudogene | 0.55 | NA | NA |
| ENSG00000204934 | ATP6V0E2-AS1 | S | lncRNA | 0.54 | 0.55 | 0.51 |
| ENSG00000235748 | SEPT14P12 | G2 | pseudogene | 0.53 | 0.19 | NA |
| ENSG00000250685 | RP11-486L19.2 | S | lncRNA | 0.53 | 0.14 | -0.20 |
| ENSG00000237870 | AC073130.1 | S | lncRNA | 0.51 | 0.15 | NA |
| ENSG00000240541 | TM4SF1-AS1 | S | lncRNA | 0.47 | -0.01 | NA |
| ENSG00000213742 | ZNF337-AS1 | S | lncRNA | 0.47 | 0.20 | NA |
| ENSG00000245213 | RP11-10K16.1 | S | lncRNA | 0.46 | 0.26 | 0.20 |
| ENSG00000250899 | RP11-253E3.3 | G1/S | lncRNA | 0.45 | 0.01 | -0.11 |
| ENSG00000214049 | UCA1 | M/G1 | lncRNA | 0.42 | 0.05 | -0.13 |
| ENSG00000281398 | SNHG4 | M/G1 | lncRNA | 0.42 | 0.42 | NA |
| ENSG00000204177 | BMS1P1 | S | pseudogene | 0.40 | 0.00 | NA |
| ENSG00000226312 | CFLAR-AS1 | S | lncRNA | 0.38 | 0.04 | NA |
| ENSG00000235897 | TM4SF19-AS1 | S | lncRNA | 0.38 | 0.49 | NA |
| ENSG00000270332 | SMC2-AS1 | G1/S | lncRNA | 0.38 | 0.44 | 0.23 |
| ENSG00000197182 | MIRLET7BHG | S | lncRNA | 0.37 | 0.21 | NA |

| | | | | | | |
|---|---|---|---|---|---|---|
| ENSG00000278864 | RP11-45M22.2 | M/G1 | TEC | 0.37 | 0.25 | -0.27 |
| ENSG00000248092 | NNT-AS1 | G1/S | lncRNA | 0.37 | 0.17 | NA |
| ENSG00000267100 | ILF3-AS1 | G1/S | lncRNA | 0.37 | 0.33 | NA |
| ENSG00000204054 | LINC00963 | S | lncRNA | 0.35 | 0.18 | NA |
| ENSG00000264538 | SUZ12P1 | G1/S | pseudogene | 0.35 | 0.35 | NA |
| ENSG00000241769 | LINC00893 | S | lncRNA | 0.34 | 0.01 | -0.59 |
| ENSG00000225138 | CTD-2228K2.7 | S | lncRNA | 0.29 | 0.00 | -0.06 |
| ENSG00000280239 | CTB-50L17.8 | G1/S | TEC | 0.28 | 0.37 | NA |
| ENSG00000214826 | DDX12P | S | pseudogene | 0.28 | 0.13 | NA |
| ENSG00000255198 | SNHG9 | G1/S | lncRNA | 0.27 | 0.43 | -0.09 |
| ENSG00000225733 | FGD5-AS1 | S | lncRNA | 0.26 | -0.01 | NA |
| ENSG00000230479 | AP000695.6 | S | lncRNA | 0.25 | 0.17 | NA |
| ENSG00000233016 | SNHG7 | M/G1 | lncRNA | 0.25 | 0.29 | 0.27 |
| ENSG00000237886 | NALT1 | S | lncRNA | 0.25 | 0.17 | NA |
| ENSG00000232445 | RP11-132A1.4 | M/G1 | lncRNA | 0.25 | 0.33 | 0.53 |
| ENSG00000274605 | RP11-12G12.7 | S | lncRNA | 0.24 | -0.18 | 0.05 |
| ENSG00000268573 | RP11-158H5.7 | S | lncRNA | 0.24 | -0.30 | NA |
| ENSG00000274173 | RP4-568C11.4 | S | lncRNA | 0.21 | 0.06 | 0.45 |
| ENSG00000197989 | SNHG12 | M/G1 | lncRNA | 0.20 | 0.13 | 0.44 |
| ENSG00000254635 | WAC-AS1 | S | lncRNA | 0.20 | -0.15 | NA |
| ENSG00000215417 | MIR17HG | M/G1 | lncRNA | 0.19 | 0.12 | -0.40 |
| ENSG00000177410 | ZFAS1 | G1/S | lncRNA | 0.18 | 0.36 | 0.43 |
| ENSG00000224032 | EPB41L4A-AS1 | M/G1 | lncRNA | 0.16 | 0.51 | 0.31 |
| ENSG00000233901 | LINC01503 | S | lncRNA | 0.16 | 0.29 | -0.17 |
| ENSG00000269893 | SNHG8 | G1/S | lncRNA | 0.15 | 0.24 | 0.38 |
| ENSG00000255717 | SNHG1 | G1/S | lncRNA | 0.14 | -0.31 | 0.06 |
| ENSG00000267080 | ASB16-AS1 | S | lncRNA | 0.13 | 0.04 | NA |
| ENSG00000259488 | RP11-154J22.1 | G1/S | lncRNA | 0.12 | -0.02 | -0.31 |
| ENSG00000154874 | CCDC144B | S | pseudogene | 0.11 | -0.28 | 0.16 |
| ENSG00000226950 | DANCR | G1/S | lncRNA | 0.11 | -0.09 | 0.07 |
| ENSG00000251562 | MALAT1 | S | lncRNA | 0.10 | 0.28 | NA |
| ENSG00000254531 | FLJ20021 | S | lncRNA | 0.08 | -0.27 | NA |
| ENSG00000272468 | RP1-86C11.7 | M/G1 | lncRNA | 0.08 | -0.22 | NA |
| ENSG00000236384 | LINC00479 | S | lncRNA | 0.07 | -0.05 | NA |
| ENSG00000254837 | AP001372.2 | S | lncRNA | 0.06 | -0.28 | 0.00 |
| ENSG00000260260 | SNHG19 | M/G1 | lncRNA | 0.06 | 0.28 | -0.01 |
| ENSG00000234327 | AC012146.7 | S | lncRNA | 0.05 | -0.21 | 0.25 |
| ENSG00000231298 | LINC00704 | S | lncRNA | 0.04 | 0.08 | NA |
| ENSG00000268798 | CTB-25B13.5 | M/G1 | lncRNA | 0.04 | -0.03 | 0.18 |
| ENSG00000268364 | SMC5-AS1 | S | lncRNA | 0.04 | -0.45 | NA |
| ENSG00000261578 | RP11-21L23.2 | G2/M | lncRNA | 0.04 | NA | NA |
| ENSG00000204860 | FAM201A | S | lncRNA | 0.03 | -0.34 | NA |
| ENSG00000180385 | EMC3-AS1 | G1/S | pseudogene | 0.00 | -0.10 | -0.18 |
| ENSG00000241015 | TPM3P9 | G1/S | pseudogene | 0.00 | 0.52 | 0.12 |
| ENSG00000236751 | LINC01186 | S | lncRNA | 0.00 | -0.18 | -0.50 |
| ENSG00000251381 | LINC00958 | G1/S | lncRNA | -0.02 | 0.23 | 0.28 |

| | | | | | | |
|---|---|---|---|---|---|---|
| ENSG00000234741 | GAS5 | M/G1 | lncRNA | -0.02 | 0.08 | -0.49 |
| ENSG00000258727 | RP11-66N24.3 | S | lncRNA | -0.05 | -0.30 | -0.07 |
| ENSG00000230937 | MIR205HG | S | lncRNA | -0.07 | 0.17 | 0.18 |
| ENSG00000279873 | LINC01126 | S | lncRNA | -0.07 | -0.02 | NA |
| ENSG00000203709 | C1orf132 | S | lncRNA | -0.08 | -0.44 | -0.36 |
| ENSG00000249042 | CTD-2015H6.3 | S | lncRNA | -0.12 | -0.04 | NA |
| ENSG00000253669 | KB-1732A1.1 | S | lncRNA | -0.13 | -0.30 | NA |
| ENSG00000237813 | AC002066.1 | S | lncRNA | -0.15 | -0.52 | NA |
| ENSG00000232995 | RGS5 | S | lncRNA | -0.17 | -0.20 | NA |
| ENSG00000187951 | ARHGAP11B | G2 | lncRNA | -0.18 | -0.29 | NA |
| ENSG00000206337 | HCP5 | G2 | lncRNA | -0.21 | -0.42 | NA |
| ENSG00000235609 | AF127936.9 | G1/S | lncRNA | -0.23 | NA | NA |
| ENSG00000260708 | CTA-29F11.1 | M/G1 | lncRNA | -0.28 | -0.19 | NA |
| ENSG00000260032 | LINC00657 | S | lncRNA | -0.29 | -0.39 | NA |
| ENSG00000263412 | RP5-890E16.2 | G1/S | lncRNA | -0.32 | -0.01 | NA |
| ENSG00000269896 | RP4-740C4.5 | S | pseudogene | -0.35 | 0.27 | 0.00 |
| ENSG00000227036 | LINC00511 | S | lncRNA | -0.38 | -0.37 | -0.16 |
| ENSG00000230844 | ZNF674-AS1 | M/G1 | lncRNA | -0.42 | -0.52 | -0.51 |
| ENSG00000203875 | SNHG5 | G1/S | lncRNA | -0.61 | -0.35 | NA |
| ENSG00000244879 | GABPB1-AS1 | S | lncRNA | -0.80 | -0.78 | NA |

TEC = To be Experimentally confirmed

**Supplementary Table S3: Info about the four lncRNA candidates.**

| Ensembl Gene ID | lncRNA names | Localization | Strand | Transcripts | Coding probability (CP*) |
|---|---|---|---|---|---|
| ENSG00000224032 | EPB41L4A-AS1<br>TIGA1 | Chr5: 112,160,526-112,164,818 | Plus | 5 | 0.1416 |
| ENSG00000177410 | ZFAS1 | Chr20: 49,278,178-49,299,600 | Plus | 14 | 0.0730 |
| ENSG00000232445 | RP11-132A1.4<br>EMS<br>AC006329.1 | Chr7: 101,308,270-101,314,800 | Plus | 2 | 0.0167 |
| ENSG00000228649 | AC005682.5<br>SNHG26 | Chr7: 22,854,126-22,872,945 | Plus | 13 | 0.0077 |

*CP values are results from the Coding Potential Assessment Tool. Human CP cutoff is 0.364. CP >= 0.364 indicates coding sequence, whereas CP < 0.364 indicates noncoding sequence (2).

**Supplementary Table S4: Primers used for RT-qPCR.** Qiagen QuantiTect primer assays (249900) were used for mRNA expression analysis and Qiagen $RT^2$ lncRNA PCR assays (330701) were used for lncRNA expression analysis.

| Gene | Primer assay | Cat.no |
|---|---|---|
| GAPDH | Hs_GAPDH_1_SG QuantiTect Primer Assay | QT00079247 |
| PCNA | Hs_PCNA_1_SG QuantiTect Primer Assay | QT00024633 |
| TOP2A | Hs_TOP2A_1_SG QuantiTect Primer Assay | QT00037632 |
| CCNB1 | Hs_CCNB1_1_SG QuantiTect Primer Assay | QT00006615 |
| CCNE2 | Hs_CCNE2_1_SG QuantiTect Primer Assay | QT00063511 |
| AC005682.5 | $RT^2$ lncRNA qPCR Assay for Human LOC101927841 | LPH07369A-200 |
| EPB41L4A-AS1 | $RT^2$ lncRNA qPCR Assay for Human EPB41L4A-AS1 | LPH04387A-200 |
| RP11-132A1.4 | $RT^2$ lncRNA qPCR Assay for Human RP11-132A1.4 | LPH09823A-200 |
| ZFAS1 | $RT^2$ lncRNA qPCR Assay for Human ZFAS1 | LPH01402A-200 |

**Supplementary Table S5: Human control qPCR primer sets from Active Motif.** Human control qPCR primer sets are designed to serve as positive or negative ChIP controls when performing ChIP with human samples.

| Primer set | Location | Cat.no | Control for |
|---|---|---|---|
| Human positive control primer set ACTB-2 | ACTB promoter | 71005 | H3K4me3 and Pol II |
| Human positive control primer set GAPDH-2 | GAPDH intron 1 | 71006 | H3K4me3 and Pol II |
| Human positive control primer set MYT1 | MYT1 promoter | 71007 | H3K27me3 |
| Human positive control primer set CCND2 | CCND2 promoter | 71008 | H3K27me3 |
| Human negative control primer set 1 | Gene desert on chr. 12 | 71001 | H3K4me3, Pol II and H3K27me3 |

**Supplementary Table S6: Antisense oligos and siRNAs used in RNA interference experiments.**

| Target lncRNA | Alias | Cat.no | Producer | Sense sequence |
|---|---|---|---|---|
| EPB41L4a-AS1 | E1 | CTM-431273 | Dharmacon | ACAGUGAGGAUGUGAAUAAUU |
| EPB41L4a-AS1 | E2 | CTM-431274 | Dharmacon | GUGAGGAUGUGAAUAAAUAUU |
| ZFAS1 | Z1 | Custom | Sigma | CUUUGAUUGAACCAGGAUG |
| ZFAS1 | Z2 | Custom | Sigma | AGUAGAAUAUAUAUAUACA |
| RP11-132A1.4 | R1 | Custom | Sigma | CACAGGAUGUGCAGAUCUC |
| RP11-132A1.4 | R2 | Custom | Sigma | GAAAUGUCAUAUAGAGAAU |
| AC005682.5 | A1 | Custom | Sigma | CUGCAUUCCUUGUCCUCAU |
| AC005682.5 | A2 | CTM-431277 | Dharmacon | GAAAGGAGGCUGAGUAAUUUU |
| AC005682.5 | ASO | 339517 LG00221049 | Qiagen | GGAAGCCACAGTAGTA |
| siRNA neg. control | Ctr | SIC001 | Sigma | |
| ASO neg. control A | Ctr | 339515 LG00000002 | Qiagen | AACACGTCTATACGC |
