## Supplementary Methods for "Joint changes in RNA, RNA polymerase II, and promoter activity through the cell cycle identify non-coding RNAs involved in proliferation"

### Supplementary method 1- RNA-seq library preparation

Briefly, 900 ng total RNA was used as starting material. The first step involved the removal of ribosomal RNA (rRNA) using biotinylated, target-specific oligos combined with Ribo-Zero rRNA removal beads. Following purification, RNA was fragmented into small pieces using divalent cations at 94°C for 4 min. First and second strand cDNAs were synthesized using random oligonucleotides and SuperScript II, followed by DNA polymerase I and RNase H. Exonuclease/polymerase was used to produce blunted overhangs. Illumina SR adapter oligonucleotides were ligated to the cDNA after 3' end adenylation. DNA fragments were enriched by 15 cycles of PCR. The libraries were purified using the AMPure XP (Beckman Coulter, Inc., Indianapolis, IN, USA), quantitated by qPCR using KAPA Library Quantification Kit (Kapa Biosystems, Inc., Wilmington, MA, USA), and validated using Agilent High Sensitivity DNA Kit on a Bioanalyzer (Agilent Technologies, Santa Clara, CA, USA). The size range of the DNA fragments was measured to be in the range of 210-850 bp and peaked around 310 bp. Prior to sequencing, the libraries were quantified (KAPA Library Quantification Kit (Illumina/ABI Prism), normalized, pooled, and size selected (Ampure XP beads). Then each pool was quantified again. The pooled purified libraries were normalized and 20 pM was subject to clustering (by a cBot Cluster Generation System on a HiSeq2500 Rapid run mode flow cell (Illumina)), according to manufacturer's instructions.

### Supplementary method 2-Chromatin Immunoprecipitation (ChIP)

For each time point in the cell cycle, ~ 3.5 million HaCaT cells were washed twice with PBS, cross-linked with 0.5% formaldehyde at room temperature for 10 min, following addition of 125 mM glycine to stop the cross-linking. Cells were washed again with PBS and scraped from the plates in PBS supplemented with 1mM EDTA and protease inhibitors. Cross-linked cells were collected by centrifugation and resuspended in 1 ml cold RIPA buffer (10 mM Tris pH 8.0, 1 mM EDTA, 140 mM NaCl, 1% Triton X-100, 0.1% SDS, 0.1% Na-Deoxycholate) supplemented with protease inhibitors and 125 mM glycine. Samples were frozen in liquid nitrogen and stored at -80°C. Later, cells were thawed and sonicated, using a model 250 Sonifier (Branson Ultrasonic Corporation), at 20% duty cycle, 2.5 output control, 1 min x 10 with 30 sec pauses in between. To further digest the DNA, cells were treated with 400 U of Micrococcal Nuclease (NEB, M02475) at 37°C for 15 min. Debris was removed by centrifuging (16.000 x g) at 4°C for 15 min and soluble chromatin supernatant equivalent to 3.5 x $10^6$ cells/IP was pre-cleared using 20 µl of a 1:1 mix of protein A and protein G Dynabeads (Invitrogen). Overnight immunoprecipitation was performed using either two µg of anti-H3K4me3 (Diagenode, C1541003-50), anti-H3K27me3 (Diagenode, C15410195), anti- Pol II (Diagenode, C15200004), or the non-specific immunoglobulin G (IgG) antibodies Mouse IgG (Diagenode, C15400001) and Rabbit IgG (Diagenode, C15410206). Antibody-protein complexes were immunoprecipitated with protein A/G Dynabeads for 3 hours at 4°C, then washed five times with RIPA buffer supplemented with protease inhibitors, once with LiCl wash buffer (250 mM LiCl, 10 mM Tris pH 8.0, 1 mM EDTA, 0.5% NP-40, 0.5% Na-Deoxycholate) supplemented with protease inhibitors, and once in TE (10 mM Tris-HCl pH 8.0, 1 mM EDTA). Immunoprecipitated DNA was reverse cross-linked by adding 1 µl RNaseA [10 mg/ml] and incubated at 37°C for 30 min. Then we added 2.5 µl 20% SDS and 5 µl Proteinase K [10mg/ml] and incubated at 55°C for 1 hour, followed by purification using QIAquick PCR purification kit (Qiagen, 28104).

**Supplementary method 3- ChIP-seq library preparation**

ChIP-seq libraries were prepared using the MicroPlex Library Preparation Kit v4 (Diagenode) in accordance with the manufacturer's protocol. In brief, ~1 ng immunoprecipitated dsDNA was used as input in the template preparation process that provides efficient end repair of the fragmented dsDNA input. Further, library synthesis was performed by ligation of MicroPlex patented stem-loop adapters. The stem-loop adaptors with blocked 5' ends were ligated to the 5' end of the genomic DNA, leaving a nick at the 3'end. Next, a library amplification step was performed, which enabled extension of the template, cleavage of the stem-loop adaptors, and amplification of the library. In the final step, the 3' ends of the genomic DNA were extended to complete library synthesis and Illumina-compatible indexes were added through a high-fidelity amplification. Prior to sequencing, the libraries were quantified using the KAPA Library Quantification Kit (Illumina/ABI Prism), normalized, pooled, and size selected (Ampure XP beads). Then each pool was quantified again. The pooled purified libraries were normalized, and 20 pM was subject to clustering (by a cBot Cluster Generation System on a HiSeq2500 HO flow cell (Illumina)), according to manufacturer's instructions.

**Supplementary method 4- Western blot analysis**

In the cell synchronization experiments, cells were washed once with cold PBS before pelleted, snap-frozen, and stored at -80°C. Total protein extracts for Pol ll analysis were prepared by resuspending cell pellets in 1 x pellet-cell-volume (PCV) of Buffer I (10 mM Tris-HCl pH 8.0, 200 mM KCl, 1 mM DTT, protease inhibitor) and 1 x PCV of Buffer II (10 mM Tris-HCl pH 8.0, 200 mM KCl, 2 mM EDTA, 40% Glycerol, 0.5% Triton X-100, 0.5% NP-40, 1 mM DTT, protease inhibitor). The mixture was gently shaken for 2 hours at 4°C, extracts were cleared by centrifugation, and protein concentrations were measured using the Bradford method (Bio-Rad). Total protein extracts for H3K4me3 and H3K27me3 analysis were isolated using Allprep DNA/RNA/protein mini Kit (Qiagen, 80004) according to the manufacturer's protocol, except that the lysis buffer in the last step was replaced with SDS buffer. The protein concentration was measured using Direct Detect assay free cards (Merc, DDAC00010-GR) and analyzed on a Direct Detect Infrared Spectrometer. Proteins were separated by electrophoresis on NuPAGE 4-12% Bis-Tris gels (Invitrogen, NP0322BOX) and blotted onto PVDF membranes by standard procedures. PVDF membrane for high molecular weight proteins (Amersham Hybond, 10600023) was used for RNA Pol II detection and PVDF membrane for low molecular weight proteins (Amersham Hybond, 10600061) was used for H3K4me3 and H3K27me3 detection. Primary antibodies for RNA Pol II (Diagenode, C15200004), H3K4me3 (Diagenode, C15410003), and H3K27me3 (Diagenode, C15410195) were diluted [1:1000] in 5% dry milk in PBS containing 0.1% Tween and membranes were incubated overnight at 4°C. Anti-beta Actin (Abcam, ab8226) [1:5000] was used as loading control. The secondary antibodies IRDye 800CW goat anti-rabbit (LI-COR Biosciences, 925-32211) and 680RD goat anti-mouse (LI-COR Biosciences, 926-68070) were used in a [1:15000] dilution and membranes were visualized using the LI-COR Biosciences detection system.
