## Supplementary figures and images for "Joint changes in RNA, RNA polymerase II, and promoter activity through the cell cycle identify non-coding RNAs involved in proliferation"

### Graphical abstract

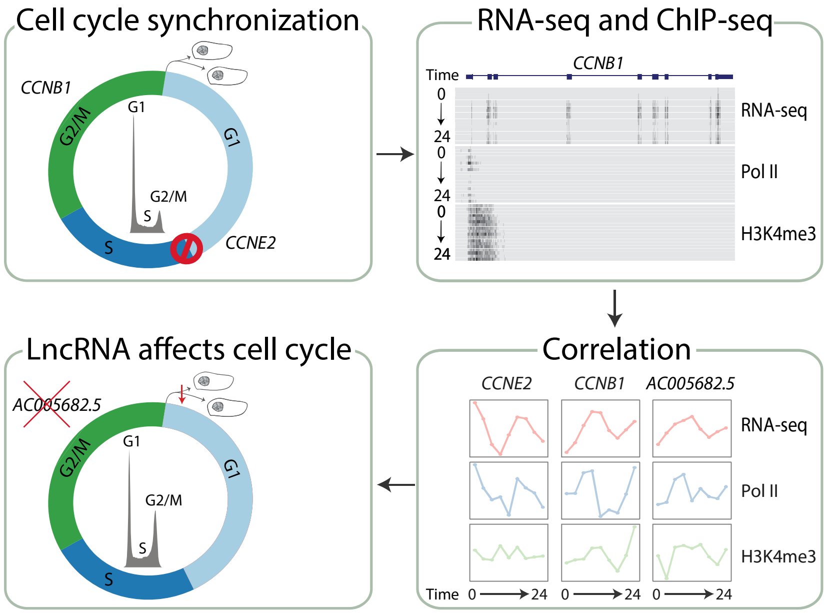
